# HERC1-mediated orphan subunit quality control maintains proteasome homeostasis and is disrupted in HERC1-associated neurodevelopmental disease

**DOI:** 10.64898/2026.11.09.765296

**Authors:** Manel Garcia-Diez, Laura Costa-Sastre, Joan Sala-Gaston, Tania López-Hernández, Juanma Ramírez, Nerea Osinalde, Jose Antonio Valer, Claudia Arnedo-Pac, Ugo Mayor, Francesc Ventura, Jose Luis Rosa

**Author notes:** Senior author (Jose Luis Rosa).

## Abstract

Biallelic hypomorphic variants in the E3 ubiquitin ligase HERC1 cause a neurological disorder clinically characterized by macrocephaly, dysmorphic facies, and psychomotor delay. Identifying the substrates of HERC1 is essential for understanding its biological functions and the mechanisms underlying disease-associated variants. To address this, we performed a quantitative proteomic analysis using biotinylated ubiquitin to identify HERC1 targets and examine their regulation in cells expressing HERC1 with or without ubiquitin ligase activity. This approach revealed a broad set of HERC1-dependent ubiquitylation substrates, including proteins from major multiprotein complexes such as the proteasome, the translation initiation machinery, and pathways involved in cell metabolism. Focusing on the proteasome, the most prominently represented complex, we identified several regulatory particle subunits, including PSMC6, PSME1, and PSME2. Specifically, within the 19S regulatory particle, we demonstrated that ubiquitylation of the PSMC6 subunit involves the formation of K48-linked polyubiquitin chains and depends on HERC1 activity. Other 19S subunits, such as PSMC2, PSMC4, and PSMC5, undergo a similar modification. Mechanistically, we show that HERC1 selectively recognizes unassembled subunits, promotes their ubiquitylation, and targets them for proteasomal degradation. Notably, in HERC1- deficient cells, disruption of this mechanism impaired proteasomal activity. At the disease level, quantitative proteomic analysis of the pathogenic HERC1 variant c.1448G>A (p.Gly483Glu) likewise identified altered ubiquitylation of proteasome subunits and components of the translation initiation machinery. Biochemical analyses further revealed impaired ubiquitylation of PSMC6 by the mutant HERC1 protein, associated with a loss of interaction between HERC1 and PSMC6. Together, these findings establish a link between HERC1-associated neurological disorders and defective ubiquitylation pathways. They elucidate the molecular mechanisms by which HERC1 regulates proteostasis and demonstrate how disruption of these processes contributes to disease pathogenesis, thereby highlighting potential avenues for therapeutic intervention.

## INTRODUCTION

Pathogenic variants in the E3 ubiquitin ligases HERC1 and HERC2 are linked to distinct neurological disorders. Biallelic hypomorphic HERC2 variants cause a neurodevelopmental syndrome clinically resembling Angelman syndrome, characterized by global developmental delay, intellectual disability, autism spectrum features, and movement abnormalities (OMIM #615516) [1–7]. Similarly, biallelic HERC1 variants result in *Macrocephaly, Dysmorphic Facies, and PsychoMotor Retardation* (MDFPMR) syndrome (OMIM #617011) [6,8]. Mouse models, most notably the *tambaleante Herc1* mutant and *Herc2* haploinsufficient mice, recapitulate key neurological and cellular phenotypes, including Purkinje cell degeneration, synaptic dysfunction, and impaired motor learning [6,9–11]. However, despite the growing genetic and phenotypic evidence linking HERC1 and HERC2 dysfunction to neurological disease, the molecular pathways connecting these ubiquitin ligases to neuronal homeostasis remain poorly understood. A major barrier to closing this gap is the limited knowledge of their physiological substrates. Identifying the proteins directly regulated by HERC1 and HERC2 will be key to defining the signaling networks they govern and to explaining how their disruption culminates in neuronal dysfunction and disease. Using a biotinylated ubiquitin (^bio^Ub) strategy combined with comparative quantitative proteomics[12], we identified a broad set of HERC1-dependent ubiquitylation targets, including numerous subunits of multimeric protein complexes. Interaction studies and functional assays further establish that HERC1 plays a central role in proteasome assembly and activity, revealing proteasome homeostasis as a conserved function of HERC family ubiquitin ligases. Moreover, quantitative proteomic profiling of a pathogenic HERC1 variant, together with complementary biochemical approaches, demonstrates that HERC1 dysfunction converges on impaired proteasome homeostasis, providing a link between HERC1-associated neurological disorders and defective ubiquitylation pathways.

## METHODS

### Cell culture and treatments

HEK-293T and MCF7 human cells were obtained and cultured as previously described[13]. Where indicated, cells were treated with MG132 (10μM) for the indicated time. Plasmid transfections were carried out using Lipofectamine LTX transfection reagent or Polyethylenimine (PEI) according to the manufacturer’s instructions. siRNA-mediated knockdown, shRNA-mediated knockdown, and lentiviral particle production and cell infection were performed as previously described[13]. MISSION shRNA clone of human HERC1 (TRCN0000235499) was purchased from Sigma-Aldrich (St. Louis, MA).

### Constructs

pBsSVD2005 (Addgene #21826) was a gift from David Ron. pMD2.G (Addgene #12259) and psPAX2 (Addgene #12260) were a gift from Didier Trono. pLKO.1 - TRC control was a gift from David Root (Addgene #10879). His-Ub and His-Ub-K48R were a gift from Tatiana Erazo. KY28 (EGFP-P2A-mCherry), EZ103 (EGFP-PSMC5-P2A-mCherry, EZ144 (EGFP-PSMC4-P2A- mCherry), EZ125 (3xFlag-PAAF1), EZ127 (3xFlag-HERC1), EZ179 (3xFlag-HERC1 ΔHECT), and EZ180 (3xFlag-HERC1 G483E) plasmids were a gift from Ramanujan S. Hegde[14]. JLR247 (EGFP-PSMC6-P2A-mCherry) was generated by *in vivo* homologous-recombination–based assembly[15], using a human fetal brain cDNA library (Stratagene, La Jolla, CA) as the source of PSMC6 and KY28 (EGFP-P2A-mCherry) as the vector backbone. pCAG-(^bio^Ub)x6-BirA was previously described[12].

### Protein extraction, SDS-PAGE, and immunoblotting

Protein extraction, SDS-PAGE, and immunoblotting were performed as previously described[13]. Antibodies used: anti-HERC1[16]; anti-Flag M2 (Sigma-Aldrich, F3165 and F7425); anti-GFP (Abcam, ab13970); anti-α-tubulin (Abcam, ab7291); anti-Ran[16]; anti-PSMC6 (ProteinTech #15839-1-AP); and anti-clathrin heavy chain (sc12734), anti-PSMC2 (sc-166972), anti-PSMC4 (sc-166115), anti-PSMC5 (sc390631), and anti-vinculin (sc73614) from Santa Cruz Biotechnology. Anti-mouse (#926-32210), anti-rabbit (#926-68071) and anti-chicken (#925- 68075) antibodies conjugated to fluorochromes from LICORbio were used as secondary antibodies when Odyssey M imager (LI-COR Biotech, Lincoln, USA) was used.

### Pull-down assay

To analyze protein-protein interactions, GFP pull-down assays were performed as previously described[14]. To analyze ubiquitylation, assays were performed as previously described[13] using the urea lysis buffer (50 mM HEPES pH 7.4, 150 mM NaCl, 8 M urea, 1% CHAPS, 5 mM imidazole and 10 mM 2-mercaptoethanol) for lysis, incubation and washing.

### Immunocytochemistry and confocal imaging

Cells were analyzed as previously described[13] and using the antibodies indicated for immunoblotting. Antibodies conjugated to Alexa-Fluor 488 or 647 (1:500; Thermo Fisher Scientific) were used as secondary antibodies.

### Quantitative proteomic analysis with the ^bio^Ub strategy

Biotin pull-downs and proteomic analysis were performed in triplicates as previously described[12,17]. The mass spectrometry results have been deposited to the ProteomeXchange Consortium[18] with the dataset identifier PXD081793.

### Functional connectivity networks

Proteins with significantly increased ubiquitylation in HERC1 WT were analyzed using STRING (v.12.0)[19].

### Proteasome activity and GFP fluorescence analysis in native PAGE

For proteasome activity, equal amounts of protein (5-20 µg) were loaded by triplicated in 96-well dark plates containing 200 μL/well of Proteasome buffer supplemented with 50μM Suc-LLVY- AMC (Sigma-Aldrich, S6510), and incubated for 60 min at 37°C in dark. After this time, fluorescence was measured using a CLARIOstar Plus microplate reader (BMG LABTECH, Ortenberg, Deutschland). See more details in[17].

For GFP fluorescence analysis in native PAGE, equal amounts of protein (5-20 µg) were loaded in 3-10% Tris-Acetate PAGE and running with native gel buffer[17]. After electrophoresis, GFP fluorescence was registered using an Odyssey M imager (LI-COR Biotech, Lincoln, USA).

### Statistical analysis

Data were analyzed using GraphPad Prism version 8.4.3 and expressed as means and standard error of the mean (± SEM) of, at least, three independent experiments. Significance was calculated by unpaired Student’s t-tests and indicated as follows: *, **, or *** for p-values of ≤ 0.05, ≤ 0.01, or ≤ 0.001, respectively.

## RESULTS

### ^Bio^Ub-driven proteomics reveals HERC1-dependent ubiquitylation networks

To uncover substrates whose ubiquitylation depends on HERC1, we employed a ^bio^Ub-based enrichment strategy combining transient expression of distinct HERC1 variants with label free quantitative proteomics[12]. This approach relies on *in vivo* biotinylation of ubiquitin, achieved by expressing the ^bio^Ub-BirA construct (Fig. 1a). The construct encodes a polyprotein consisting of six ubiquitin units, each engineered at the N-terminus with a short biotin-acceptor peptide, followed by the promiscuous *E. coli* BirA biotin ligase fused at the C-terminus. After expression, endogenous deubiquitinases process the precursor, releasing individual ubiquitin molecules that are subsequently biotinylated by BirA. To distinguish HERC1-dependent ubiquitylation events, together with the ^bio^Ub-BirA construct, we co-expressed either wild-type HERC1 (HERC1 WT) or a catalytically inactive mutant lacking the catalytic domain (HERC1 ΔHECT). Biotinylated, and hence, ubiquitylated proteins were captured using avidin affinity and identified by mass spectrometry. Quantitative comparison of samples expressing HERC1 WT versus HERC1 ΔHECT enabled the selective identification of proteins whose ubiquitylation depends on HERC1 activity (Fig. 1a).

**Figure 1.**
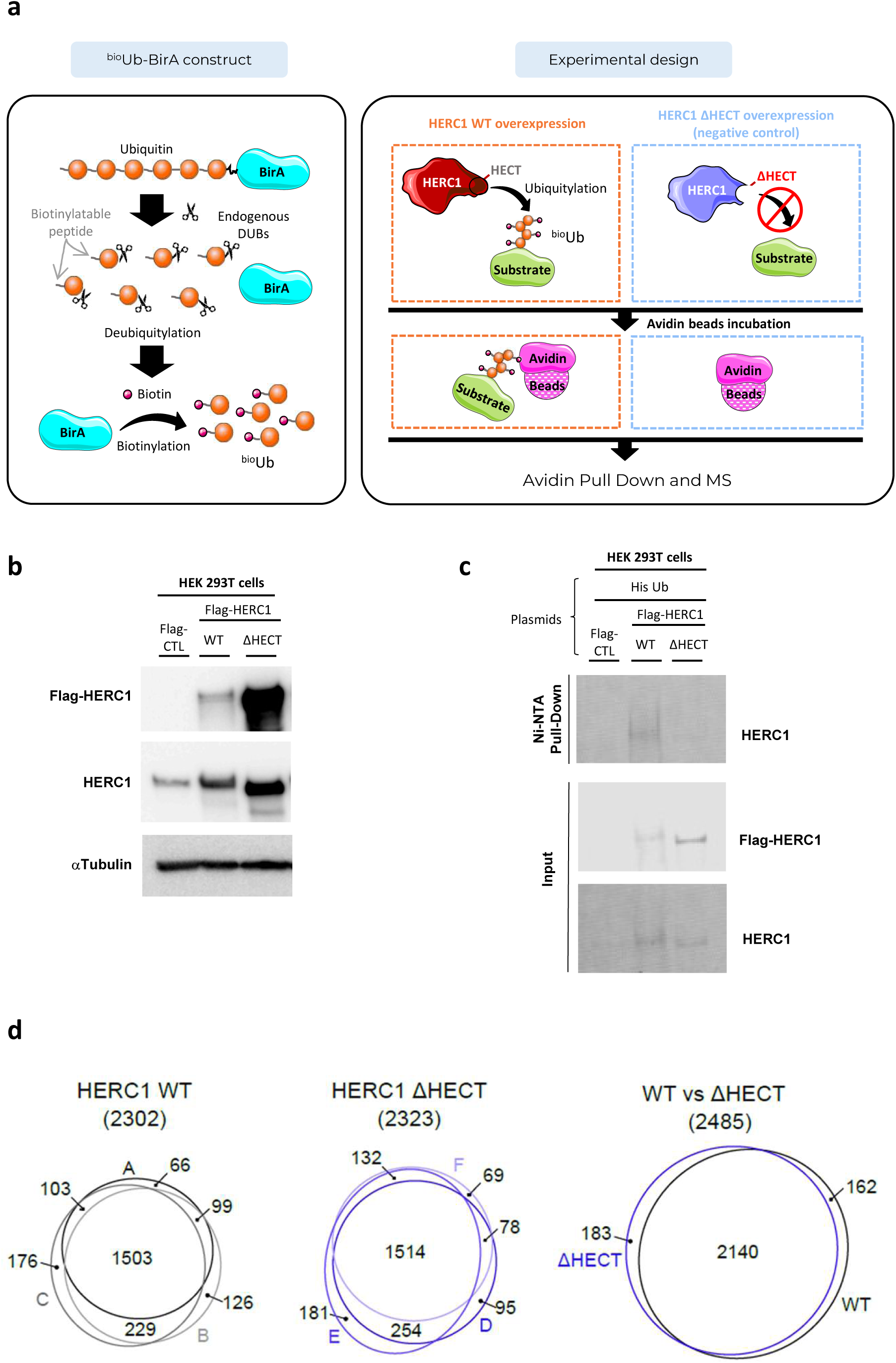
^Bio^Ub-based proteomics identifies protein ubiquitylation regulated by HERC1. (a) The ^bio^Ub strategy is based on the *in vivo* biotinylation of ubiquitin (Ub) achieved by transfecting the ^bio^Ub-BirA construct, and combined with the overexpression of the wild-type HERC1 (HERC1 WT) or a catalytically inactive mutant lacking the catalytic domain (HERC1 ΔHECT). Ubiquitylated proteins are isolated with avidin beads and identified by mass spectrometry (MS). (b) HEK-293T cells were transfected with a negative control plasmid (Flag-CTL), a plasmid encoding HERC1 WT protein fused with Flag epitope (Flag-HERC1 WT) or a plasmid encoding a catalytically inactive form (Flag-HERC1 ΔHECT). The indicated proteins were analyzed by immunoblot. (c) HERC1 regulates its own ubiquitylation. HEK-293T cells were transfected with the indicated plasmids. After 48 hours, cells were incubated for 6h with MG132 (10 µM). Ubiquitylated proteins were purified using a Ni-NTA-agarose resin. Inputs and pull-downs were analyzed by immunoblotting with antibodies against the indicated proteins. Representative results are shown. (d) Reproducibility of the mass spectrometry analysis. Overlap between proteins identified in HERC1 WT (depicted in black) and HERC1 ΔHECT samples (depicted in blue). Venn diagrams showing the overlap of the proteins identified among the three replicas of each condition, as well as between both conditions are shown.

Lysates from transfected HEK-293T cells were first examined by western blotting, which confirmed robust expression of both HERC1 constructs, with the catalytically inactive variant accumulating to higher levels than the WT (Fig. 1b). This observation suggested that HERC1 may undergo autoregulation through ubiquitylation-dependent proteasomal degradation. To test this possibility, we performed an *in vivo* ubiquitylation assay using His-tagged ubiquitin. Following pull-down of His-ubiquitylated proteins, samples were analyzed by SDS-PAGE followed by immunoblotting. A pronounced high-molecular-weight smear, indicative of HERC1 polyubiquitylation, was readily detected in cells expressing HERC1 WT, whereas the signal was markedly reduced in the HERC1 ΔHECT mutant. These results demonstrate that HERC1 undergoes autoubiquitylation, supporting a model in which the E3 ligase activity promotes its own turnover (Fig. 1c).

Triplicate lysates from HEK-293T cells co-expressing the ^bio^Ub-BirA construct and either HERC1 WT and HERC1 ΔHECT were subjected to ^bio^Ub -based enrichment of ubiquitinated proteins followed by label-free quantitative proteomic analysis. A high reproducibility across the biological replicas was observed (Fig. 1d) (Suppl. Fig. 1). A total of 2485 proteins were identified, of which 2140 were found both in WT and ΔHECT samples (Fig. 1d). Of these, 48 proteins exhibited higher ubiquitylation in cells expressing HERC1 WT than in cells expressing HERC1 ΔHECT, whereas 51 showed lower ubiquitylation. (Fig. 2a, Table 1 and Suppl. Table 1). To characterize proteins exhibiting increased HERC1-dependent ubiquitylation, we used STRING (Search Tool for the Retrieval of Interacting Genes/Proteins) to identify functional protein association networks and enriched biological processes. The analysis highlighted six connectivity clusters: proteasome, translation preinitiation and initiation, glutamine metabolism, amino sugar biosynthesis, and ribonucleoprotein complex biogenesis (Fig. 2b,c). Ranking these clusters by enrichment score highlighted the proteasome complex as one of the most prominent functional modules (Fig. 2c).

**Figure 2.**
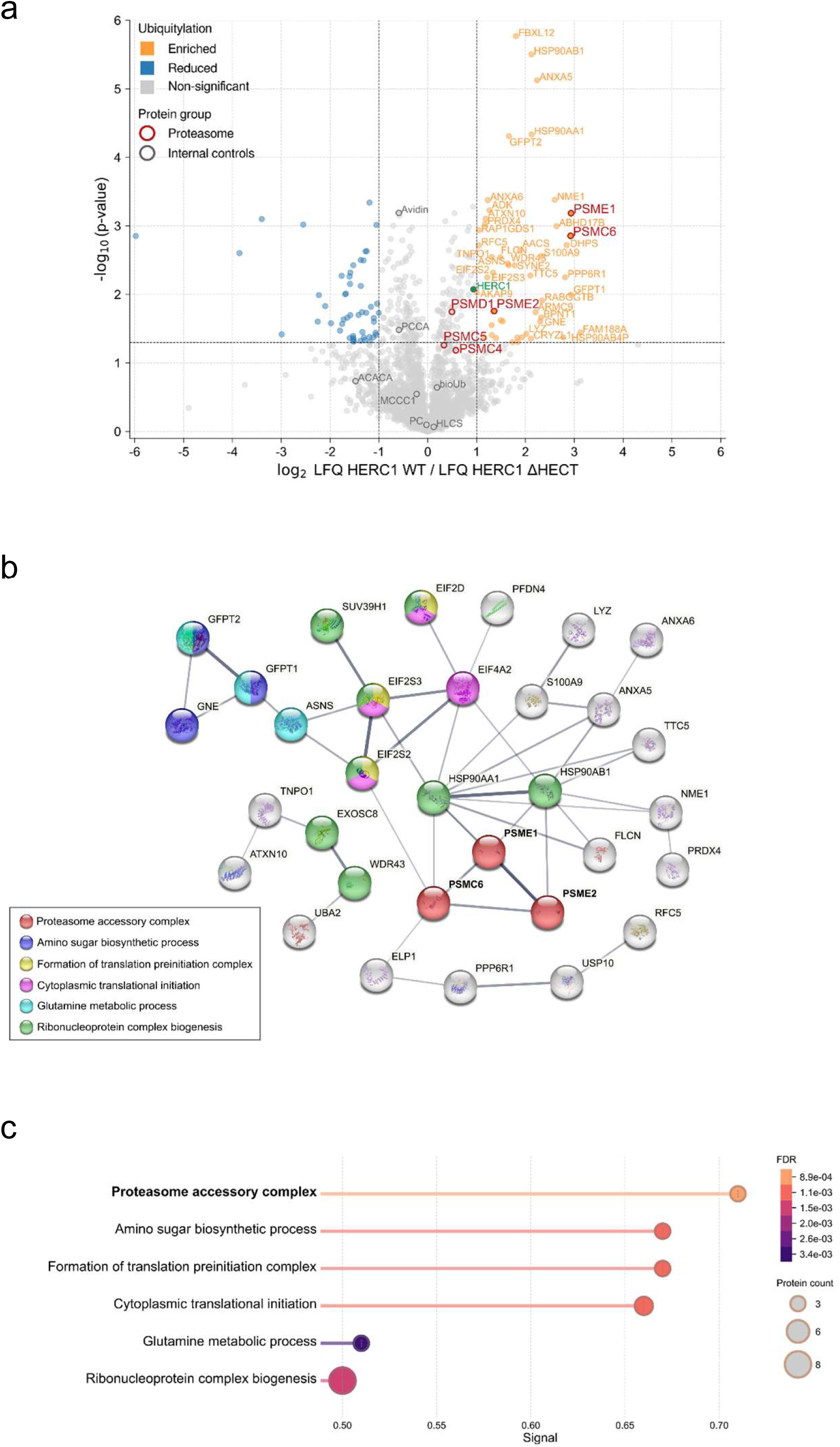
Identification of ubiquitylated proteins regulated by HERC1 and their functional connectivity. (a) Volcano plot shows differentially ubiquitylated proteins upon HERC1 WT overexpression relative to HERC1 ΔHECT. Abundance of each individual protein was determined by their label free quantification (LFQ) intensities. The LFQ WT/ΔHECT ratios (log_2_ scale) and the t-test p-values (−log_10_ scale) are displayed in the X and Y axis, respectively. The threshold for statistical significance (p-value < 0.05) is indicated with a horizontal grey dashed line, while vertical grey dashed lines depict a > 1 log_2_ fold increase or decrease of the ubiquitylated levels upon HERC1 WT overexpression. Proteins that pass these thresholds, and fulfill the quality control criteria, and show a significant increase or reduction of their ubiquitylation levels upon HERC1 WT overexpression are shown in orange and blue, respectively (log_2_ LFQ WT/LFQ ΔHECT > 1 marked in orange; log_2_ LFQ WT/LFQ ΔHECT < -1 marked in blue). HERC1 is shown in green, and endogenously biotinylated proteins (ACACA, MCCC1, PC, PCCA, HLCS), ^bio^Ub, and avidin are shown in grey. Proteasome-related proteins are shown in red. (b) Functional protein association network using STRING shows functional connectivity of ubiquitylated proteins regulated by HERC1. Grey line thickness indicates the confidence of the interaction; proteins without any interaction have been removed from the network display. (c) Functional enrichment analysis of the protein clusters detailed in (b).

**Table 1.** Ubiquitylated proteins up-regulated by HERC1.

| Gene name | Protein description | p-value (-log<sub>10</sub>) | WT/ΔHECT ratio (log<sub>2</sub>) |
|---|---|---|---|
| FBXL12 | F-box/LRR-repeat protein 12 | 5.77 | 1.80 |
| HSP90AB1 | Heat shock protein HSP 90-beta; Putative heat shock protein HSP 90-beta-3 | 5.50 | 2.12 |
| ANXA5 | Annexin A5 | 5.13 | 2.24 |
| HSP90AA1 | Heat shock protein HSP 90-alpha | 4.33 | 2.12 |
| GFPT2 | Glutamine--fructose-6-phosphate aminotransferase [isomerizing] 2 | 4.31 | 1.66 |
| NME1 | Nucleoside diphosphate kinase A; Nucleoside diphosphate kinase B | 3.38 | 2.60 |
| ANXA6 | Annexin A6 | 3.38 | 1.23 |
| ADK | Adenosine kinase | 3.23 | 1.26 |
| PSME1 | Proteasome activator complex subunit 1 | 3.18 | 2.93 |
| ATXN10 | Ataxin-10 | 3.10 | 1.19 |
| PRDX4 | Peroxiredoxin-4 | 3.02 | 1.17 |
| ABHD17B | Alpha/beta hydrolase domain-containing protein 17B | 3.00 | 2.64 |
| RAP1GDS1 | Rap1 GTPase-GDP dissociation stimulator 1 | 2.94 | 1.04 |
| PSMC6 | 26S protease regulatory subunit 10B | 2.86 | 2.92 |
| DHPS | Deoxyhypusine synthase | 2.72 | 2.84 |
| RFC5 | Replication factor C subunit 5 | 2.71 | 1.04 |
| AACS | Acetoacetyl-CoA synthetase | 2.64 | 1.89 |
| S100A9 | Protein S100-A9 | 2.56 | 2.32 |
| TNPO1 | Transportin-1 | 2.54 | 1.31 |
| FLCN | Folliculin | 2.54 | 1.49 |
| ASNS | Asparagine synthetase [glutamine-hydrolyzing] | 2.45 | 1.64 |
| WDR43 | WD repeat-containing protein 43 | 2.43 | 1.66 |
| SYNE2 | Nesprin-2 | 2.42 | 1.78 |
| EIF2S2 | Eukaryotic translation initiation factor 2 subunit 2 | 2.32 | 1.34 |
| TTC5 | Tetratricopeptide repeat protein 5 | 2.27 | 2.10 |
| EIF2S3 | Eukaryotic translation initiation factor 2 subunit 3; Putative eukaryotic translation initiation factor 2 subunit 3-like protein | 2.25 | 1.22 |
| PPP6R1 | Serine/threonine-protein phosphatase 6 regulatory subunit 1 | 2.25 | 2.81 |
| AKAP9 | A-kinase anchor protein 9 | 2.03 | 1.01 |
| GFPT1 | Glutamine--fructose-6-phosphate aminotransferase [isomerizing] 1 | 1.99 | 2.93 |
| RABGGTB | Geranylgeranyl transferase type-2 subunit beta | 1.91 | 2.34 |
| PSME2 | Proteasome activator complex subunit 2 | 1.76 | 1.36 |
| ARMC9 | LisH domain-containing protein ARMC9 | 1.74 | 2.21 |
| BPNT1 | 3(2),5-bisphosphate nucleotidase 1 | 1.66 | 2.32 |
| IKBKAP (ELP1) | Elongator complex protein 1 | 1.62 | 1.49 |
| VPS25 | Vacuolar protein-sorting-associated protein 25 | 1.61 | 1.54 |
| GNE | Bifunctional UDP-N-acetylglucosamine 2-epimerase/N-acetylmannosamine kinase; UDP-N-acetylglucosamine 2-epimerase (hydrolyzing); N-acetylmannosamine kinase | 1.60 | 2.30 |
| EXOSC8 | Exosome complex component RRP43 | 1.55 | 1.31 |
| FAM188A (MINDY3 ) | Protein FAM188A | 1.44 | 3.12 |
| LYZ | Lysozyme C | 1.42 | 2.02 |
| SUV39H1 | Histone-lysine N-methyltransferase SUV39H1 | 1.41 | 1.31 |
| HSP90AB4P | Putative heat shock protein HSP 90-beta 4 | 1.37 | 2.77 |
| EIF4A2 | Eukaryotic initiation factor 4A-II; Eukaryotic initiation factor 4A-II, N-terminally processed | 1.37 | 1.93 |
| UBA2 | SUMO-activating enzyme subunit 2 | 1.37 | 1.83 |
| VPS33B | Vacuolar protein sorting-associated protein 33B | 1.37 | 1.16 |
| EIF2D | Eukaryotic translation initiation factor 2D | 1.36 | 1.40 |
| CRYZL1 | Quinone oxidoreductase-like protein 1 | 1.36 | 2.11 |
| USP10 | Ubiquitin carboxyl-terminal hydrolase 10 | 1.31 | 1.84 |
| PFDN4 | Prefoldin subunit 4 | 1.30 | 1.74 |

### HERC1 regulates the ubiquitylation of subunits of the proteasome regulatory particle

The proteasome is the major ATP-dependent protease complex that mediates the selective degradation of intracellular proteins conjugated to polyubiquitin chains, typically linked through Lys48 (K48) residues. Through the ubiquitin–proteasome system, cells tightly regulate protein turnover, eliminate misfolded or damaged proteins, and control the abundance of key regulatory proteins involved in processes such as cell cycle progression, signal transduction, and antigen presentation, thereby maintaining proteostasis and cellular homeostasis[20,21]. In mammalian cells, the predominant proteasome species is the 30S/26S proteasome, which consists of the 20S core particle (CP) and the 19S regulatory particle (RP)[22,23]. Whereas the 20S CP harbors the proteolytic active sites, the 19S RP mediates substrate recognition, deubiquitylation, unfolding, and translocation into the CP.

The 19S RP is organized into two major subcomplexes: the base and the lid. The base contains a hetero-hexameric ring of AAA^+^-ATPases (PSMC1–6), which drive ATP-dependent substrate unfolding and gate opening of the 20S CP, together with includes several non-ATPase subunits including PSMD1, PSMD2, PSMD4, and ADRM1. Among these, PSMD2, PSMD4, and ADRM1 serve as ubiquitin receptors that recognize ubiquitylated substrates, while PSMD1 contributes to structural stabilization of the complex. The lid comprises nine non-ATPase subunits (PSMD3, PSMD6–8, PSMD11-14, and SEM1) responsible for substrate recruitment, deubiquitylation, and further stabilization of the regulatory particle. Assembly of the base and lid occurs through independent pathways and requires dedicated chaperones, including PSMD5, PSMD9, PSMD10, and PAAF1, to ensure the correct incorporation of subunits into the functional 19S RP[22,23].

Our proteomic analysis identified the ATPase subunit PSMC6, a core component of the 19S regulatory particle, as a protein exhibiting increased ubiquitylation in the presence of HERC1 (Fig. 2). This finding suggests that HERC1 may regulate proteasome assembly by modulating the turnover or incorporation of specific 19S RP subunits. Moreover, PSME1 and PSME2, which together form the PA28 (11S) proteasome activator complex, also exhibited increased ubiquitylation in the presence of HERC1 WT. Given the established role of PA28 in antigen processing and MHC class I antigen presentation [24,25], these results point to HERC1 as a potential regulator of alternative proteasome activators associated with immunoproteasome dynamics. Collectively, our findings identify suggest that HERC1 may contribute to the regulation of proteasome assembly, activity, and functional specialization.

### HERC1 interacts with unassembled 19S regulatory particle subunits and regulates their stability

To investigate the molecular mechanisms underlying HERC1-dependent changes in ubiquitylation of 19S assembly proteins identified in our proteomic analysis, we first focused on PSMC6. We demonstrated that PSMC6 ubiquitylation depends on HERC1 catalytic activity (Fig. 3a). Proteasome inhibition with MG132 increased PSMC6 ubiquitylation (Fig. 3b). Analysis of the ubiquitin chains using a K48R ubiquitin mutant indicated that PSMC6 is predominantly modified by K48-linked polyubiquitin chains (Fig. 3c and Suppl. Fig. 2), consistent with targeting for proteasomal degradation. Confocal microscopy revealed colocalization of HERC1 and PSMC6, which was enhanced upon proteasome inhibition (Fig. 3d). Pull-down assays confirmed the interaction between HERC1 and PSMC6 and further demonstrated that, in contrast to the HERC1– PSMC5 interaction[14], binding of HERC1 and PSMC6 occurs independently of the chaperone PAAF1 (Fig. 3e). Given that nascent PSMC6 subunits have a markedly shorter half-life than assembled ones[14], GFP–PSMC6 predominantly represents unassembled subunits, whereas endogenous PSMC6 is largely incorporated into mature proteasomes. Consistent with this model, siRNA-mediated depletion of HERC1 selectively increased the stability of GFP–PSMC6 (Fig. 3f) but had no effect on endogenous PSMC6 or GFP alone (Fig. 3f and Suppl. Fig. 3). Taken together, these findings support a role for HERC1 in the quality control of proteasome assembly promoting the ubiquitylation and proteasomal turnover of unassembled PSMC6.

**Figure 3.**
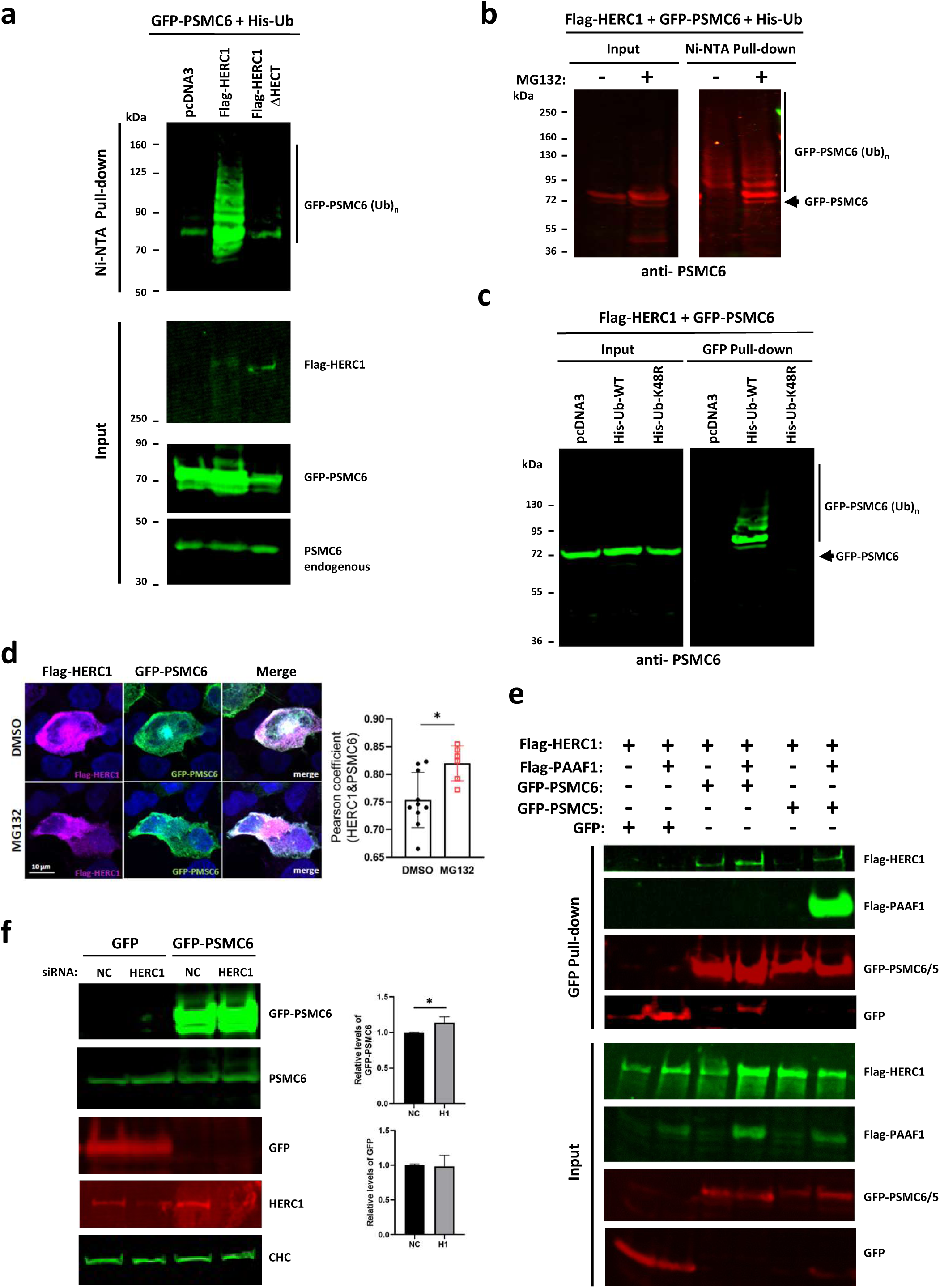
HERC1 regulates ubiquitylation and stability of unassembled PSMC6. (a) HEK- 293T cells were transfected with His-Ub, GFP-PSMC6, and either a control plasmid (Flag-CTL) or the wild-type HERC1 (Flag-HERC1) or the catalytically inactive form (Flag-HERC1 ΔHECT). After 48h, the cells were treated with proteasome inhibitor MG132 (10 μM) for 4h. Next, cell lysates were subjected to pull-down using Ni-NTA agarose. Inputs and proteins retained in the resin were analyzed by immunoblotting with antibodies against the indicated proteins. (b) HEK- 293T cells were treated and analyzed as in (a). (c) HEK-293T cells were transfected with Flag- HERC1, GFP-PSMC6, and either a control plasmid (pcDNA3), the wild-type His-Ub (WT) or the mutant His-Ub K48R. After 48h, the cells were treated with proteasome inhibitor MG132 (10 μM) for 4h. Next, cell lysates were subjected to pull-down using GFP-trap resin. Inputs and proteins retained in the resin were analyzed by immunoblotting with antibodies against PSMC6. (d) (Left) Representative confocal images (maximum intensity projections) of HEK-293T cells transfected with Flag-HERC1 and GFP-PSMC6, and incubated with either DMSO or MG132 (10 µM for 4 h), followed by immunostaining with antibodies against Flag and GFP. Scale bars, 10 μm. (Right) Quantification of co-localization between Flag-HERC1 and GFP-PSMC6, expressed as Pearson’s correlation coefficient. Data represent mean ± SEM from 54 images of DMSO-treated cells and 31 images of MG132-treated cells across 3 independent experiments. *p < 0.05, unpaired Student’s t- test. (e) HEK-293T cells were transfected with the indicated constructs. After 48h, cell lysates were subjected to pull-down using GFP-trap resin. Inputs and proteins retained in the resin were analyzed by immunoblotting with antibodies against the indicated proteins. (f) HEK-293T cells were transfected with non-targeting control (NC) or HERC1 (H1) siRNAs. After 24h, these cells were transfected again with GFP or GFP-PSMC6. 48 h later, cell lysates were analyzed by immunoblotting with antibodies against the indicated proteins. GFP and GFP-PSMC6 protein levels were quantified and normalized based on clathrin heavy chain (CHC) protein levels. The plot represents the mean ± standard error of the mean. Representative results are shown from experiments repeated three times. Significance levels: * p ≤ 0.05.

Having identified the 19S regulatory particle subunit PSMC6 as a candidate HERC1-regulated proteasome subunit, we next assessed whether additional 19S ATPase subunits are similarly regulated. To this end, we examined the ubiquitylation status of PSMC2, PSMC4 and PSMC5. Ubiquitylation assays showed that the ubiquitylation of all three subunits depends on HERC1 (Fig. 4a). Proteasome inhibition with MG132 resulted in the accumulation of ubiquitylated PSMC2, PSMC4 and PSMC5, indicating that their stability depends on proteasome activity (Fig. 4b). Analysis using a K48R ubiquitin mutant further indicated that these proteins are predominantly modified with K48-linked polyubiquitin chains (Fig. 4c), consistent with targeting for proteasomal degradation. Pull-down experiments demonstrated the interactions between HERC1 and these PSMC subunits (Fig. 4d). These experiments also demonstrated the presence of PAAF1 in these complexes. Because PAAF1 dissociates from PSMC subunits upon completion of proteasome assembly[26], its association with PSMC proteins suggests that HERC1 preferentially recognizes unassembled proteasomal subunits. Accordingly, prolonged retention of PAAF1 on PSMC subunits, indicative of defective proteasome assembly, may facilitate their recognition by HERC1, leading to their ubiquitylation and subsequent proteasomal degradation. Consistent with these findings, depletion of HERC1 increased the stability of overexpressed (and therefore predominantly unassembled PSMC subunits) while having no detectable effect on endogenous, assembled subunits (Fig. 4e). Given that nascent, unassembled proteasomal subunits exhibit a considerably shorter half-life than their assembled counterparts[14], these results highlight an important role for HERC1 as a quality-control factor for multiple 19S regulatory particle subunits by promoting ubiquitylation and subsequent proteasomal degradation of unassembled 19S ATPase subunits.

**Figure 4.**
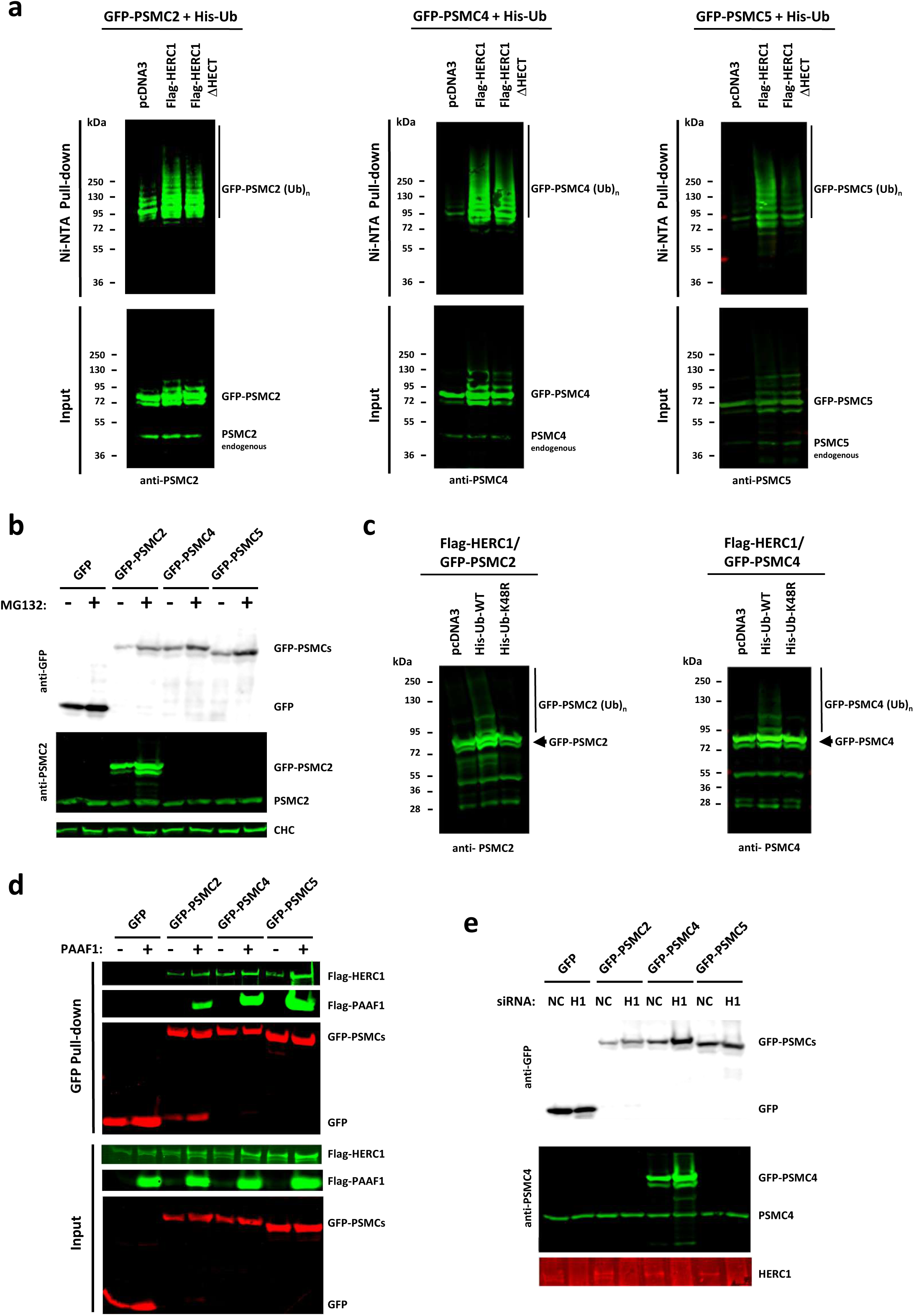
HERC1 interacts with unassembled 19S regulatory particle subunits and regulates their stability. (a) HEK-293T cells were transfected with His-Ub, the indicated GFP-PSMC (GFP- PSMC2, 4 or 5), and either a control plasmid (Flag-CTL) or the wild-type HERC1 (Flag-HERC1) or the catalytically inactive form (Flag-HERC1 ΔHECT). After 48h, the cells were treated with proteasome inhibitor MG132 (10 mM) for 4h. Next, cell lysates were subjected to pull-down using Ni-NTA agarose. Inputs and proteins retained in the resin were analyzed by immunoblotting with antibodies against the indicated proteins. (b) HEK-293T cells were treated as in (a) and lysates were analyzed by immunoblotting with antibodies against the indicated proteins. (c) HEK-293T cells were transfected with Flag-HERC1, the indicated GFP-PSMC, and either a control plasmid (pcDNA3), the wild-type His-Ub (WT) or the mutant His-Ub K48R. After 48h, the cells were treated with proteasome inhibitor MG132 (10 μM) for 4h. Next, cell lysates were analyzed by immunoblotting with antibodies against the indicated PSMC subunit. (d) HEK-293T cells were transfected with the indicated constructs. After 48h, cell lysates were subjected to pull-down using GFP-trap resin. Inputs and proteins retained in the resin were analyzed by immunoblotting with antibodies against the indicated proteins. (e) HEK-293T cells were transfected with non-targeting control (NC) or HERC1 (H1) siRNAs. After 24h, these cells were transfected again with GFP or the indicated GFP-PSMC. 48 h later, cell lysates were analyzed by immunoblotting with antibodies against the indicated proteins.

### HERC1 regulates proteasome activity

The identification of multiple 19S regulatory particle subunits as HERC1-regulated proteins suggested that HERC1 may play an important role in proteasome function. To address this possibility, we measured proteasomal activity using the fluorogenic substrate Suc-LLVY-AMC (Fig. 5a). Chronic depletion of HERC1 in HEK-293T cells (Fig. 5b) resulted in a significant reduction in proteasome activity (Fig. 5c). We hypothesized that this decrease reflects defective proteasome assembly caused by the accumulation of unassembled regulatory particle subunits. To test this possibility, cells expressing GFP–PSMC6 were analyzed by native PAGE, which preserves the integrity of proteasome complexes[17]. GFP alone served as a negative control. As expected, HERC1-depleted cells exhibited a marked accumulation of unassembled GFP–PSMC6 (blue arrow in Fig. 5d), indicating impaired clearance of excess subunits. Notably, the detection of GFP– PSMC6, but not GFP alone, within 30S/26S proteasome complexes demonstrates that GFP– PSMC6 can be incorporated into assembled proteasomes (Fig. 5d, lanes 1–2). More importantly, HERC1 depletion led to the accumulation of multiple GFP–PSMC6-containing assembly intermediates (Fig. 5d, lane 4), consistent with impaired progression of the proteasome assembly pathway. Similar results were obtained with GFP–PSMC5 (Suppl. Fig. 4). Collectively, these findings demonstrate that HERC1-mediated quality control is required for efficient proteasome assembly. In the absence of HERC1, unassembled subunits and assembly intermediates accumulate, leading to defective proteasome biogenesis and reduced proteasomal activity.

**Figure 5.**
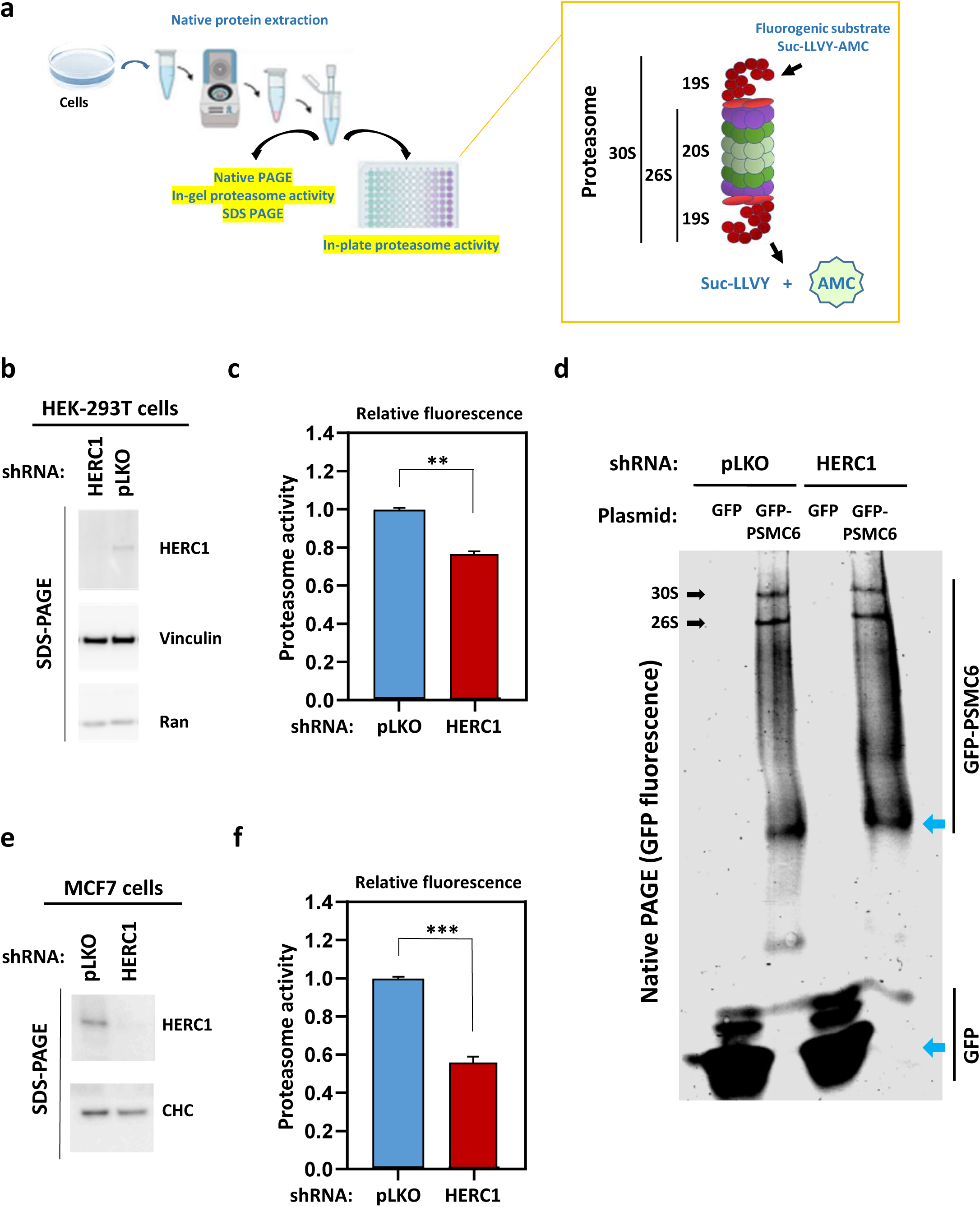
HERC1 regulates proteasome activity. (a) Strategy to analyze proteasome activity using the fluorogenic substrate Suc-LLVY-AMC. (b) Stable HEK-293T cell lines expressing a HERC1 shRNA and a non-targeting shRNA as control (pLKO) were generated. Lysates were analyzed by SDS-PAGE and immunoblotting with antibodies against the indicated proteins. (c) Proteasome activity was measured in stable HEK-293T cell lines expressing a HERC1 shRNA and a non-targeting shRNA (pLKO). Cells were lysed under native conditions. After centrifugation, supernatants were incubated with the proteasome substrate on a plate and their fluorescence measured by triplicate. Fluorescence was normalized by protein concentration and expressed relative to the control. Plots represent mean ± standard error of the mean. Experiment was repeated three times. Significance levels: ** p ≤ 0.01. (d) Stable HEK-293T cell lines expressing a HERC1 shRNA and a non-targeting shRNA (pLKO) were transfected with GFP or GFP-PSMC6. After 48h, supernatants, prepared as in (c), were analyzed in native PAGE gels. After electrophoresis, GFP fluorescence was measured. 30S and 26S structures are indicated with black arrows. Unassembled forms of GFP and GFP-PSMC6 are indicated with blue arrows. Experiment was repeated three times. (e) Stable MCF7 cell lines expressing a HERC1 shRNA and a non-targeting shRNA as control (pLKO) were generated. Lysates were analyzed by immunoblotting as in (b). (f) Proteasome activity was measured in MCF7 cell lines as in (c). Significance levels: *** p ≤ 0.001.

The effect of HERC1 depletion on proteasome activity was not restricted to HEK-293T cells but was also observed in MCF7 cells, where loss of HERC1 caused an even greater reduction in proteasomal activity (Fig. 5e, f). The increased sensitivity of MCF7 cells may be due to their aneuploid nature, which can disrupt the stoichiometric balance of proteasome subunits required for efficient complex assembly[14]. Together, these findings establish HERC1 as a critical regulator of proteasome assembly and activity across different cell types.

### HERC1-dependent ubiquitylation networks associated with a pathogenic HERC1 variant

The *tambaleante* mouse represents a valuable model for studying HERC1-associated neurological disorders, as the G483E mutation leads to defects in growth, learning, and motor coordination[6,10]. To identify changes in ubiquitylation associated with this pathogenic HERC1 variant, we performed a ^bio^Ub-based proteomic analysis analogous to that described in Fig. 1a, comparing cells expressing wild-type HERC1 (WT) with those expressing the G483E mutant (TBL). A high reproducibility across the biological replicas was observed (Fig. 6a; Suppl. Fig. 5). In total, 2,447 proteins were identified, of which 2,098 were shared between WT and TBL samples (Fig. 6a). Among these, 75 proteins exhibited differential ubiquitylation, with 37 showing increased and 38 showing decreased ubiquitylation in HERC1 WT relative to the TBL variant (Fig. 6b; Suppl. Table 2; Suppl. Table 3).

**Figure 6.**
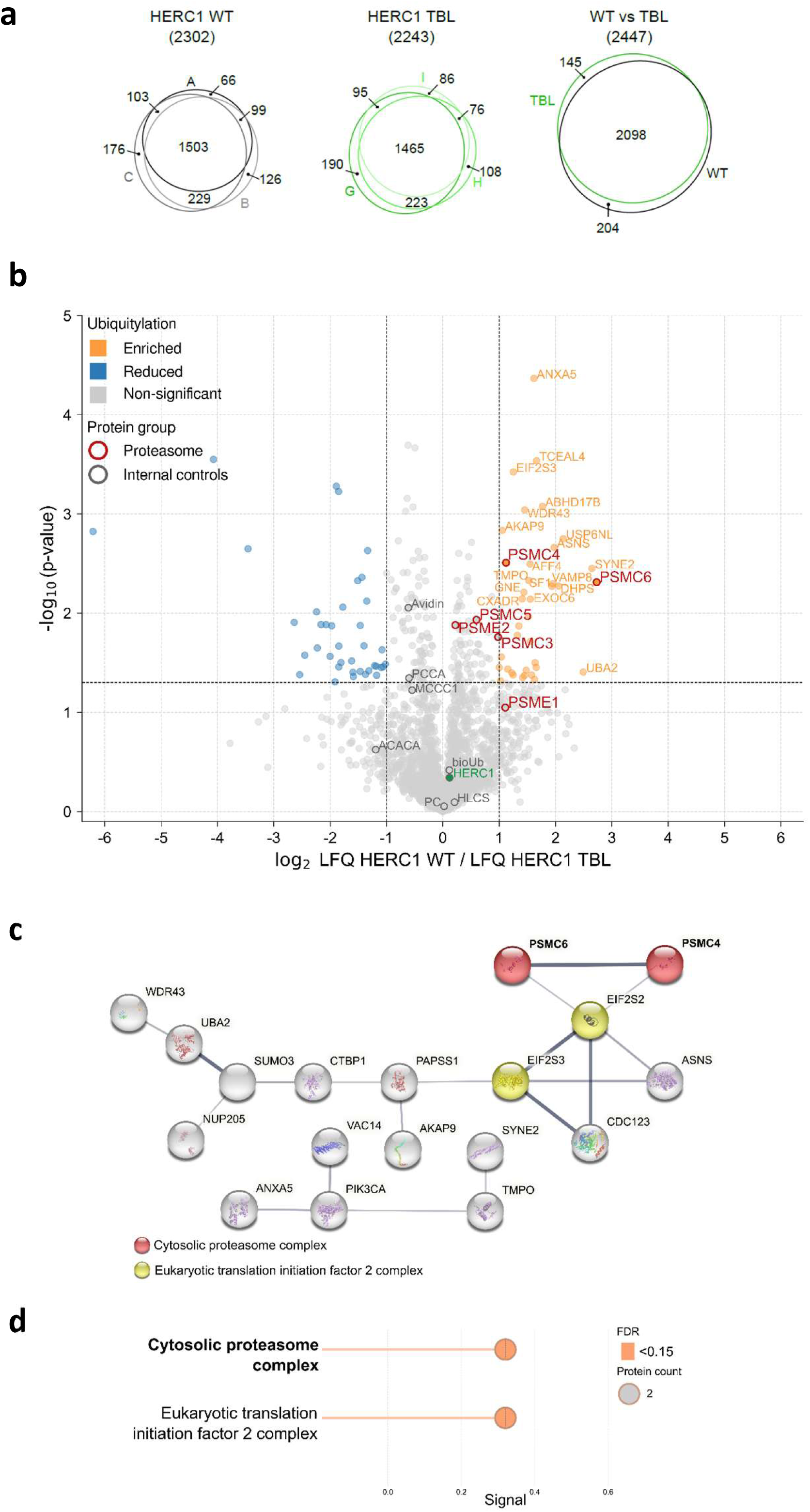
^Bio^Ub-based proteomics identifies HERC1-dependent ubiquitylation networks associated with the pathogenic HERC1 TBL variant. (a) Similar ^bio^Ub strategy described in Fig. 1 but using the pathogenic HERC1 TBL variant. Reproducibility of the mass spectrometry analysis. Overlap between proteins identified in HERC1 WT (depicted in black) and HERC1 TBL samples (depicted in green). Venn diagrams showing the overlap of the proteins identified among the three replicas of each condition, as well as between both conditions are shown. (b) Volcano plot was performed as described in Fig. 2, and shows differentially ubiquitylated proteins upon HERC1 WT overexpression relative to HERC1 TBL. (c) Functional protein association network using STRING shows functional connectivity of ubiquitylated proteins regulated by HERC1. Grey line thickness indicates the confidence of the interaction; proteins without any interaction have been removed from the network display. (d) Functional enrichment analysis of the protein clusters detailed in (c).

To investigate the functional relationship among proteins exhibiting increased ubiquitylation in HERC1 WT-expressing cells, we performed STRING network analysis. Two highly connected functional clusters emerged, corresponding to the proteasome complex and the translation initiation machinery (Fig. 6c,d). Notably, these clusters are consistent with those identified in the comparison between HERC1 WT and the catalytically inactive ΔHECT mutant (Fig. 2), reinforcing the robustness and reproducibility of the proteomic dataset. Within the proteasome cluster, PSMC4 and PSMC6 were again identified as proteins displaying increased ubiquitylation in the presence of wild-type HERC1, further implicating HERC1 in the regulation of proteasome homeostasis.

### A pathogenic HERC1 variant disrupts binding and selective ubiquitylation of unassembled PSMC6

Unlike the catalytically inactive HERC1 ΔHECT mutant, the TBL-associated HERC1 G483E variant retains an intact HECT catalytic domain. Therefore, the reduced ubiquitylation of 19S regulatory particle subunits observed in cells expressing the TBL variant is unlikely to attribute to loss of intrinsic ligase activity (Fig. 1). To gain mechanistic insight, we examined the impact of the G483E mutation on PSMC6 ubiquitylation using His–ubiquitin pull-down assays. Compared with WT HERC1, the TBL variant exhibited a reduced ability to promote the ubiquitylation of ectopically expressed GFP–PSMC6 (Fig. 7a). Consistent with our previous data, these findings suggest that the TBL variant selectively impairs the ubiquitylation of unassembled PSMC6.

**Figure 7.**
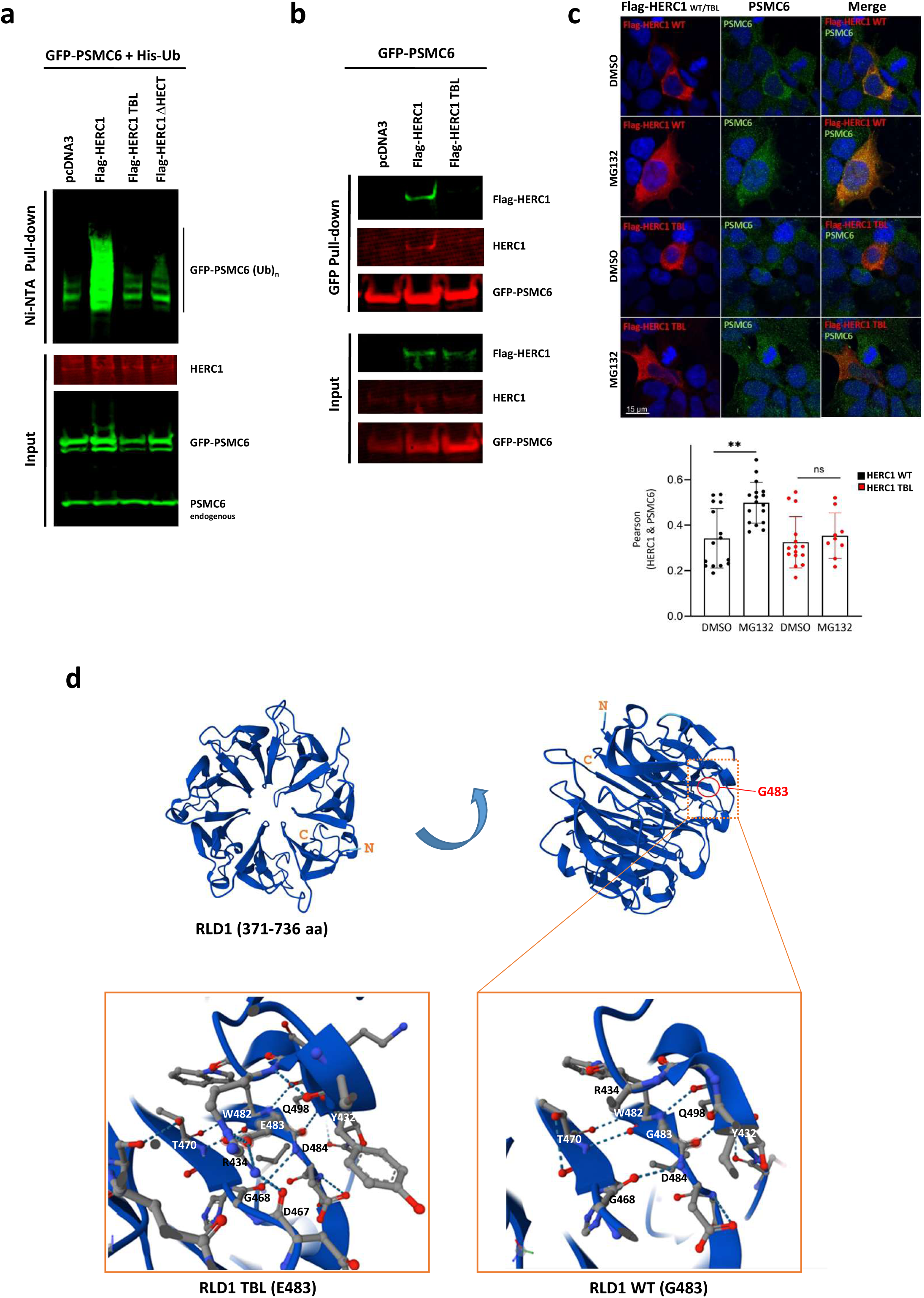
Pathogenic HERC1 TBL disrupts binding and ubiquitylation of unassembled PSMC6. (a) HEK-293T cells were transfected with His-Ub, GFP-PSMC6, and either a control plasmid (Flag-CTL) or the wild-type HERC1 (Flag-HERC1) or the catalytically inactive form (Flag-HERC1 ΔHECT) or the TBL-associated variant (Flag-HERC1TBL). After 48h, the cells were treated with proteasome inhibitor MG132 (10 μM) for 4h. Next, cell lysates were subjected to pull-down using Ni-NTA agarose. Inputs and proteins retained in the resin were analyzed by immunoblotting with antibodies against the indicated proteins. (b) HEK-293T cells were transfected with GFP-PSMC6, and either a control plasmid (pcDNA3), the wild-type HERC1 (Flag-HERC1) or the TBL variant (Flag-HERC1 TBL). After 48h, the cells were treated with proteasome inhibitor MG132 (10 μM) for 4h. Next, cell lysates were subjected to pull-down using GFP-trap resin. Inputs and proteins retained in the resin were analyzed by immunoblotting with antibodies against the indicated proteins. (c) Representative confocal images of HEK-293T cells transfected with either Flag-HERC1 WT or the TBL variant and treated with either DMSO or MG132. Cells were immunostained with anti-PSMC6 and anti-Flag antibodies. Scale bars, 15 μm. Quantification of co-localization between Flag-HERC1 WT or Flag-HERC1 TBL and endogenous PSMC6, measured using Pearson’s correlation coefficient. Data represent mean ± SEM. Significance levels: ** p ≤ 0.01; ns, not significant; one-way ANOVA followed by Tukey’s multiple-comparisons test. (d) Structural analysis of the RLD1 domain in wild-type and TBL- associated HERC1. (Top left) AlphaFold 3 prediction [34] of the seven-bladed β-propeller structure of the human HERC1 RLD1 domain (amino acid residues 371–736; UniProt: Q15751). The N- and C-terminal residues are highlighted in orange. (Right) Turn and enlarged view of the region surrounding Gly483 in the wild-type protein, showing a network of five hydrogen bonds that stabilize the local structure: two between G483 and Q498, two between W482 and T470, and one between D484 and G468. (Bottom left) Enlarged view of the corresponding region in the pathogenic G483E (TBL) variant. In addition to the five hydrogen bonds observed in the wild-type structure, three additional hydrogen bonds are predicted to form: two between E483 and R434 and one between D467 and R434. These interactions result from the recruitment of Arg434 to stabilize the negatively charged Glu483 side chain.

One possible explanation is that the G483E mutation compromises the interaction between HERC1 and PSMC6. To test this hypothesis, we performed pull-down assays and detected a reduced association between GFP–PSMC6 and the TBL variant compared with wild-type HERC1 (Fig. 7b). In agreement with these findings, confocal microscopy revealed that proteasome inhibition did not enhance the colocalization of HERC1 TBL and PSMC6, in contrast to the increased colocalization observed with HERC1 WT (Fig. 7c). Together, these results suggest that the pathogenic G483E mutation impairs substrate recognition rather than catalytic activity, thereby reducing the ubiquitylation of unassembled PSMC6. Collectively, these findings support a model in which efficient recognition of assembly intermediates is required for HERC1-mediated quality control of proteasome subunits.

## DISCUSSION

The experimental approach of combining the ^bio^Ub methodology with overexpression of the active (WT) and inactive (ΔHECT) forms of HERC1, and subsequent quantitative proteomic analysis of ubiquitylated proteins, has allowed us to identify 99 proteins regulated by ubiquitylation by this ubiquitin ligase. All these proteins represent previously unrecognized targets of HERC1-dependent ubiquitylation, and their characterization provides new insight into the molecular mechanisms regulating these targets. Functional enrichment analysis of the potential substrates revealed six major cellular processes controlled by HERC1: proteasome function, translation preinitiation and initiation, glutamine metabolism, amino sugar biosynthesis, and ribonucleoprotein complex biogenesis. Notably, two of these pathways (proteasome function and translation initiation) were also identified in analyses using a pathogenic HERC1 variant, thereby reinforcing the central role of HERC1 in regulating these processes.

Our work lays the foundation for future investigations into previously unrecognized cellular processes regulated by HERC1-dependent ubiquitylation, including eukaryotic translation initiation. Notably, we identify four translation initiation factors (EIF2S2, EIF2S3, EIF4A2, and EIF2D) as potential HERC1 substrates (Table 1), pointing to a broader role for this ligase in the control of protein synthesis. EIF4A paralogs are ATP-dependent RNA helicases that function within the eIF4F complex to promote translation initiation by unwinding secondary structures in the 5′ untranslated regions of target mRNAs, thereby facilitating ribosome scanning [27]. More recently, eIF4A2 has been shown to play a distinct role in regulating the biogenesis of 18S rRNA and the 40S ribosomal subunit [28]. EIF2S2 (eIF2β) and EIF2S3 (eIF2γ) are subunits of the heterotrimeric eIF2 complex (eIF2αβγ), which, together with GTP and initiator methionine tRNA (Met-tRNAᵢ), forms the eIF2 ternary complex (eIF2–GTP–Met-tRNAᵢ). This complex is essential for translation initiation, as it delivers the initiator tRNA to the 40S ribosomal subunit and serves as a key regulatory node linking cellular stress to global protein synthesis[27]. In contrast, EIF2D is a noncanonical translation initiation factor that promotes initiator tRNA recruitment to the 40S ribosomal subunit independently of eIF2, particularly under conditions where ternary complex formation is impaired, such as during cellular stress[29]. The identification of EIF2D therefore suggests that HERC1 may regulate not only canonical translation initiation but also alternative, eIF2-independent pathways that sustain protein synthesis under stress conditions.

The proteasome emerges as the most prominent biological process regulated by the ubiquitin ligase activity of HERC1. Functional analyses identified up to six subunits of the proteasome regulatory particle as potential targets. Four of these (PSMC2, PSMC4, PSMC5, and PSMC6) are AAA⁺ ATPases that assemble into the heterohexameric ring forming the base of the 19S regulatory particle. The remaining two proteins, PSME1 and PSME2 (also known as PA28α and PA28β, respectively), constitute an ATP-independent 11S regulatory complex which, together with the catalytic core, forms the immunoproteasome, primarily involved in the processing of MHC class I peptides [24,25]. Proteasome proteolytic activity depends on its abundance, which is tightly regulated at the level of subunit assembly[23]. Although further structural studies will be required to fully elucidate how HERC1-mediated ubiquitylation of individual subunits impacts regulatory particle assembly, the analyses presented here provide a framework for understanding the underlying molecular mechanisms. Unassembled proteasomal subunits exhibit markedly shorter half-lives than their assembled counterparts[14]. Our data demonstrate that HERC1 specifically regulates the stability of these unassembled subunits. Mechanistically, HERC1 interacts with unassembled PSMC proteins. In some cases, including PSMC2, PSMC4, and PSMC5, PAAF1 is also associated with the complex. These interactions likely facilitate the ubiquitylation of unassembled PSMC subunits, targeting them for proteasomal degradation. The chaperone PAAF1, which is absent from the fully assembled proteasome, plays a critical role in regulatory particle assembly. It binds to the carboxy-terminal domain of unassembled PSMC4 and PSMC5, promoting and coordinating their incorporation into the complex [26,30,31]. In the case of PSMC5, our findings are consistent with previous observations supporting this mechanism[14]. Importantly, our study provides the first evidence that defects in HERC1-dependent regulation of proteasome subunit assembly impair proteasome activity. Notably, a recent study identified HERC2, the other member of the Large HERC family, as a quality control factor in proteasome assembly, acting through the ubiquitylation of unassembled 19S regulatory particle subunits[17]. Taken together, these findings identify proteasome homeostasis as a shared functional hallmark of Large HERC ubiquitin ligases, establishing this family as a central regulator of proteasome assembly and function.

Assembly quality control pathways are essential guardians of proteostasis and cell viability. Their importance is particularly evident in long-lived, post-mitotic cells such as neurons, which depend on these mechanisms to prevent age-related proteotoxic stress [32]. Consistent with this, neurological dysfunction frequently arises upon perturbation of Large HERC proteins [6]. In humans, biallelic variants in *HERC1* cause neurodevelopmental disorders characterized by macrocephaly, dysmorphic facial features, and psychomotor delay [6,8]. Likewise, biallelic hypomorphic variants of *HERC2* are associated with neurodevelopmental disorders resembling Angelman syndrome, including global developmental delay, intellectual disability, autism, and movement disorders [1–7].

In mice, haploinsufficiency of *HERC2* results in impaired proteostasis in Purkinje cells, leading to neuronal loss and deficits in motor coordination [11]. Severe neurodegeneration is also observed in the *tambaleante* (TBL) mouse model, which expresses the pathogenic *HERC1* G483E variant [10]. Together, these observations underscore the critical role of Large HERC ubiquitin ligases in maintaining neuronal proteostasis. The present work provides mechanistic insight into how disruption of HERC1 function contributes to neurological disease.

Proteomic analysis of the TBL-associated HERC1 variant identified the proteasome as one of the most significantly affected biological processes. In particular, several subunits of the 19S regulatory particle, including PSMC4 and PSMC6, displayed reduced ubiquitylation in cells expressing the TBL variant. Subsequent functional studies focused on PSMC6 confirmed a marked decrease in its ubiquitylation. Importantly, because the catalytic HECT domain remains intact in the TBL variant, this defect is unlikely to result from a loss of intrinsic E3 ligase activity. Rather, our biochemical analyses indicate that the reduced interaction between HERC1 G483E and PSMC6 compromises substrate recognition, thereby decreasing the efficiency of PSMC6 ubiquitylation. At the structural level, the G483E substitution affects a highly conserved glycine residue located within RCC1-like domain 1 (RLD1) of HERC1[10]. RLD domains constitute one of the defining structural features of the HERC family of ubiquitin ligases[33] and adopt a characteristic seven- bladed β-propeller architecture (Fig. 7d). Each blade is composed of four antiparallel β-strands arranged radially around a central axis, generating a stable circular scaffold that frequently serves as a platform for protein–protein interactions. AlphaFold-based structural prediction[34] of the HERC1 RLD1 domain suggests that, in the wild-type protein, the region surrounding Gly483 is stabilized by a network of five hydrogen bonds: two between G483 and Q498, two between W482 and T470, and one between D484 and G468 (Fig. 7d, RLD1 WT). In the pathogenic G483E variant, these five hydrogen bonds are retained, but three additional hydrogen bonds are predicted to form: two between E483 and R434 and one between D467 and R434 (Fig. 7d, RLD1 TBL). These additional interactions arise from the recruitment of Arg434 to stabilize the negatively charged glutamate residue and are predicted to alter the local conformation and dynamic properties of the domain. Although the overall β-propeller architecture is preserved, these localized structural rearrangements are likely sufficient to impair substrate binding. Consistent with this interpretation, the G483E mutation reduces the interaction of HERC1 with PSMC6 and may consequently diminish its ubiquitylation. Previous reports showing altered interactions of HERC1 G483E with the PSMC5–PAAF1 complex[14] and with clathrin[35] further support the notion that the pathogenic mutation primarily disrupts protein–protein interactions mediated by the RLD1 domain rather than the catalytic activity of HERC1 itself. Collectively, these findings provide a mechanistic explanation for how the TBL-associated HERC1 variant compromises proteasome regulation through defective substrate recognition and ubiquitylation.

Our findings establish a direct link between HERC1-associated neurological disorders and impaired proteasome activity, providing mechanistic insight into how mutations in RLD domains disrupt proteostasis and drive disease pathology. These results also have potential implications for preventive strategies. Early genetic screening for HERC1 variants may enable the identification of individuals at risk. In such cases, it will be important to explore whether the resulting cellular defects can be mitigated or prevented by restoring proteasome function and rebalancing proteostasis.

## AVAILABILITY OF DATA AND MATERIALS

The mass spectrometry results have been deposited to the ProteomeXchange Consortium via the PRIDE [18] partner repository with the dataset identifier PXD081793. Original images are available on raw image file. Any additional information reported in this paper is available from the lead contact upon request.

## AUTHOR CONTRIBUTIONS

M.G.D., J.S.G., L.C.S., T.L.H., J.R., U.M., F.V., and J.L.R. conceived and designed the experiments. Performed the experiments: M.G.D., J.S.G., L.C.S., T.L.H., J.R., N.O., J.A.V., C.A., U.M., and J.L.R. All authors analyzed the data. M.G.D., J.S.G., and J.L.R. wrote the manuscript. All authors contributed to the manuscript. J.L.R. supervised the project.

## ACKNOWLEDGEMENTS

We thank E. Adanero, A. Muhaisen, E. Castaño, B. Torrejon, the Biology-Bellvitge Unit from Scientific and Technological Centers (CCiTUB), Universitat de Barcelona for technical assistance. We sincerely thank Drs. R.S. Hegde, D. Root, Tatiana Erazo, and D. Chan for kindly providing us with plasmids. Manel Garcia-Diaz is a recipient of a STEP-FI fellowship from the Generalitat de Catalunya. Laura Costa-Sastre is a recipient of a PRE-DOCS UB fellowship from the Universitat de Barcelona. This research was supported by grant PID2023-150549OB-I00 funded by MICIU/AEI/ 10.13039/501100011033 and by “ERDF A way of making Europe”, “ERDF/EU”. In addition, this work is funded by Generalitat de Catalunya (2021 SGR 00316) and based upon work from COST Action ProteoCure CA20113, supported by COST (European Cooperation in Science and Technology).

---

**Supplementary Figure 1.**
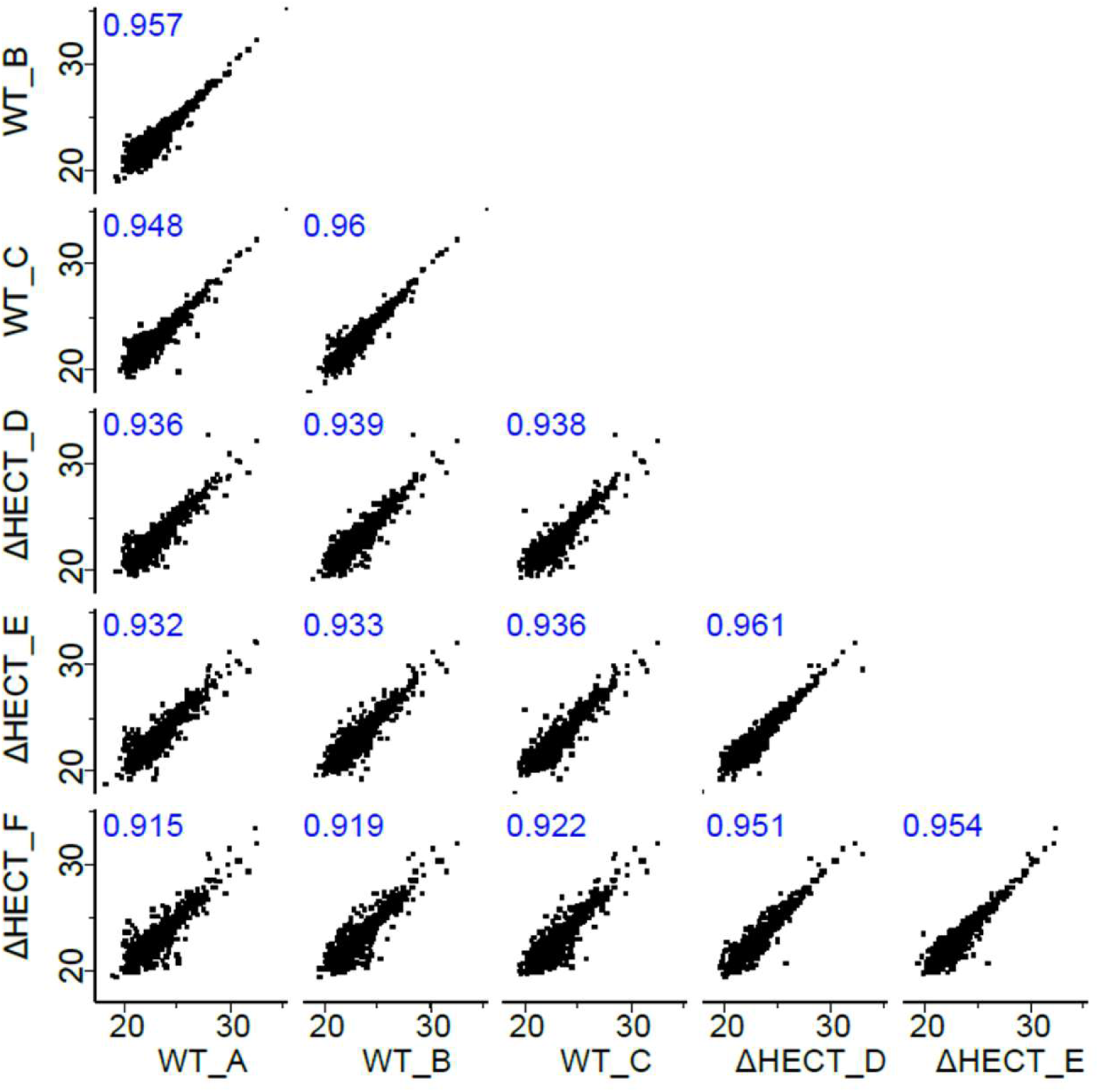
Reproducibility of the mass spectrometry analysis. Correlation graphs of label free quantification (LFQ) intensities (in log_2_ scale) of proteins identified across the different samples. Three replicas of each condition. Pearson’s correlation coefficient is shown in blue.

**Supplementary Figure 2.**
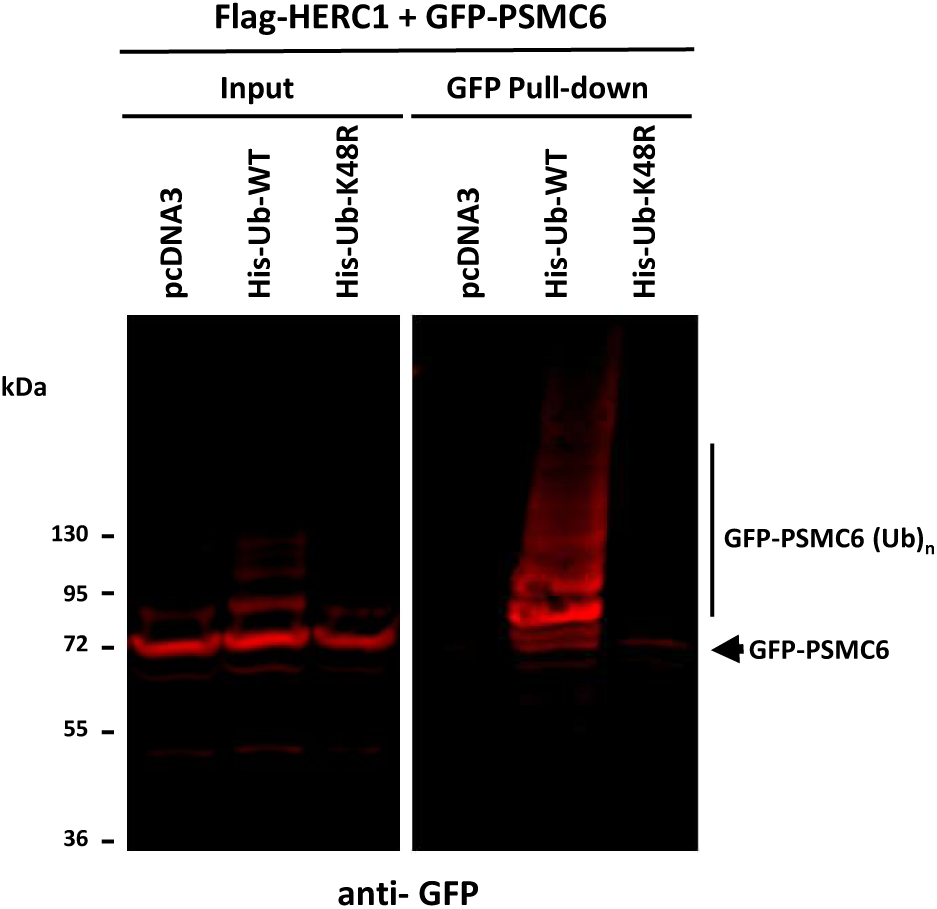
GFP-PSMC6 ubiquitylation was associated with K48-linked polyubiquitin chains. HEK-293T cells were transfected with Flag- HERC1, GFP-PSMC6, and either a control plasmid (pcDNA3), the wild-type His- Ub (WT) or the mutant His-Ub K48R. After 48h, the cells were treated with proteasome inhibitor MG132 (10 µM) for 4h. Next, cell lysates were subjected to pull-down using GFP-trap resin. Inputs and proteins retained in the resin were analyzed by immunoblotting with antibodies against GFP.

**Supplementary Figure 3.**
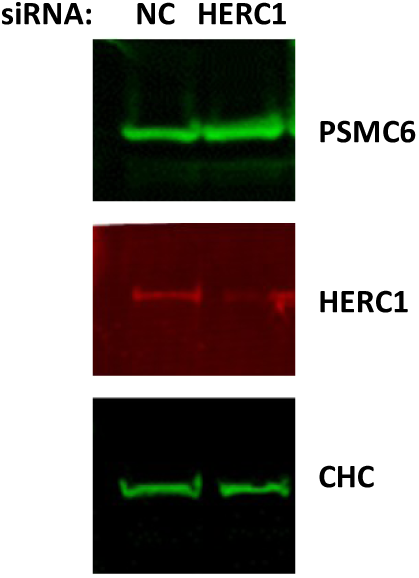
siRNA-mediated depletion of HERC1 had no effect on endogenous PSMC6. HEK-293T cells were transfected with non-targeting control (NC) or HERC1 siRNAs. After 72h, cell lysates were analyzed by immunoblotting with antibodies against the indicated proteins.

**Supplementary Figure 4.**
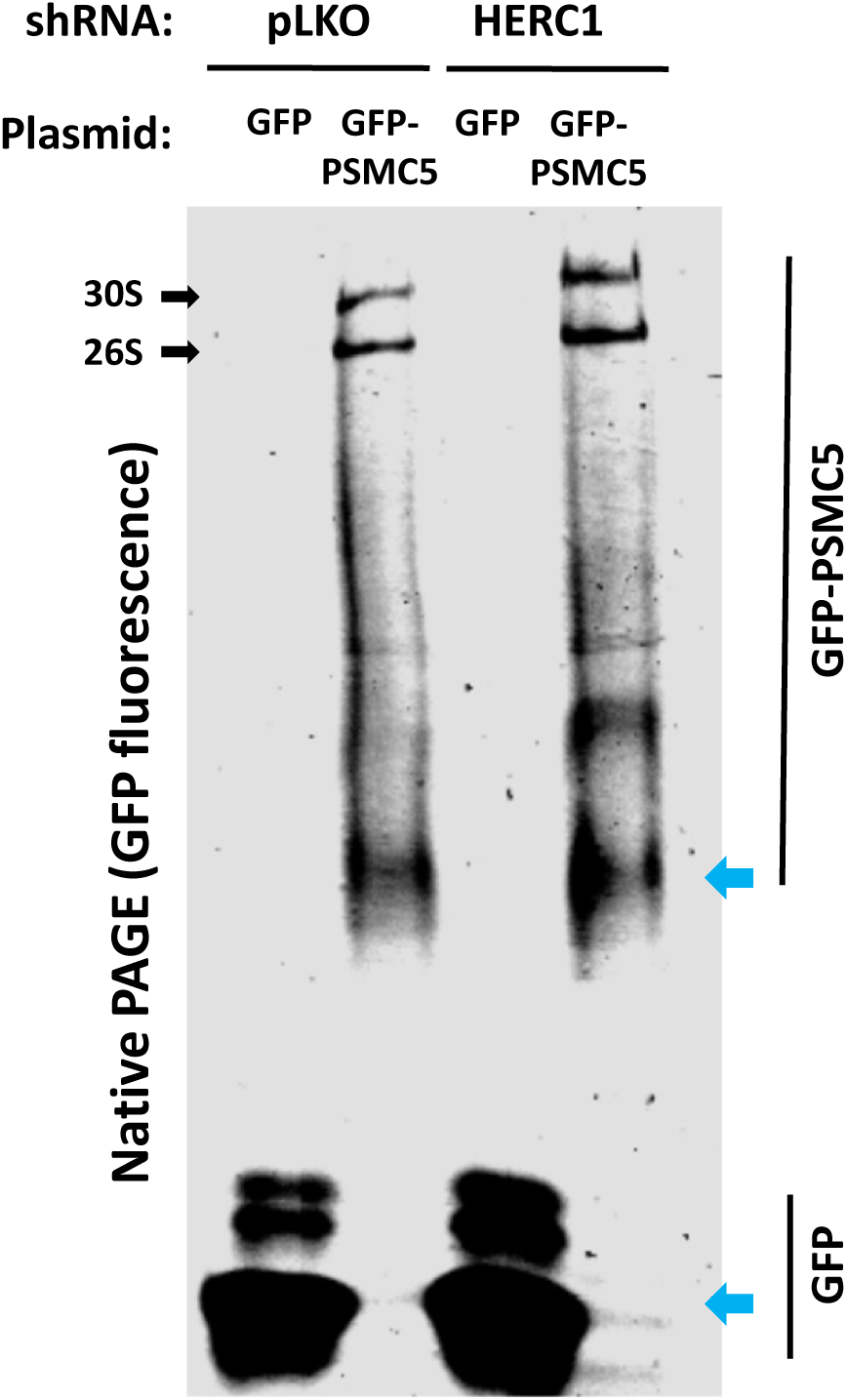
HERC1 depletion led to the accumulation of multiple GFP–PSMC5-containing assembly intermediates. Stable HEK- 293T cell lines expressing a HERC1 shRNA and a non-targeting shRNA (pLKO) were transfected with GFP or GFP-PSMC5. After 48h, supernatants, prepared as in Fig. 5, were analyzed in native PAGE gels. After electrophoresis, GFP fluorescence was measured. 30S and 26S structures are indicated with black arrows. Unassembled forms of GFP and GFP- PSMC6 are indicated with blue arrows.

**Supplementary Figure 5.**
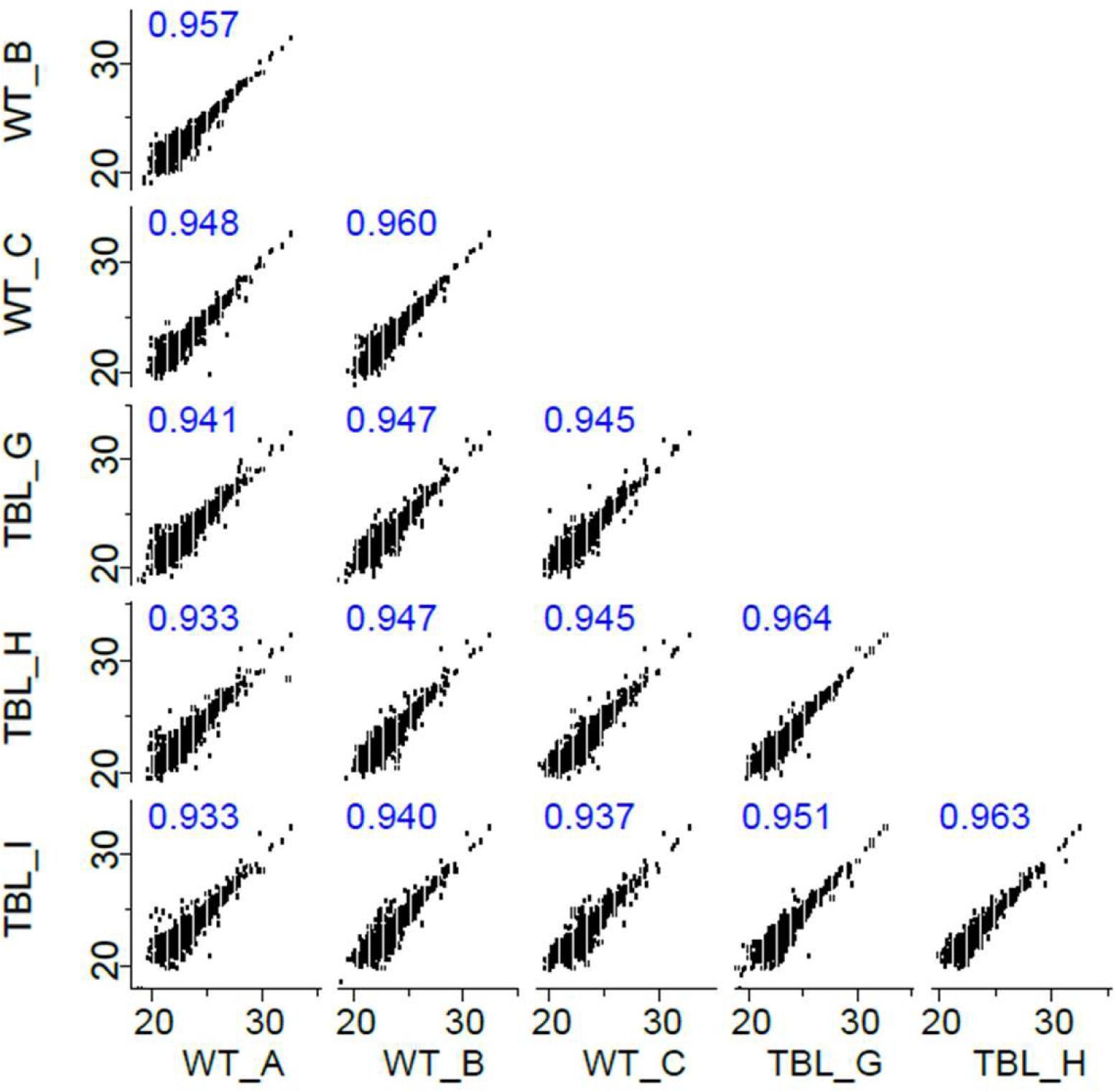
Reproducibility of the mass spectrometry analysis. Correlation graphs of label free quantification (LFQ) intensities (in log_2_ scale) of proteins identified across the different samples. Three replicas of each condition. Pearson’s correlation coefficient is shown in blue.

**Supplementary table 1.** Ubiquitylated proteins down-regulated by HERC1.

| Gene name | Protein description | p-value (-log<sub>10</sub>) | WT/ΔHECT ratio (log<sub>2</sub>) |
|---|---|---|---|
| YARS (YARS1) | Tyrosine--tRNA ligase, cytoplasmic; Tyrosine--tRNA ligase, cytoplasmic, N-terminally processed | 3.34 | -1.19 |
| TANK | TRAF family member-associated NF-kappa-B activator | 3.10 | -3.40 |
| EHD1 | EH domain-containing protein 1 | 3.02 | -2.55 |
| GIGYF2 | PERQ amino acid-rich with GYF domain-containing protein 2 | 3.01 | -1.05 |
| CS | Citrate synthase, mitochondrial | 2.85 | -5.98 |
| MPRIP | Myosin phosphatase Rho-interacting protein | 2.63 | -1.25 |
| QSER1 | Glutamine and serine-rich protein 1 | 2.63 | -1.27 |
| TIMM13 | Mitochondrial import inner membrane translocase subunit Tim13 | 2.60 | -3.86 |
| OFD1 | Oral-facial-digital syndrome 1 protein | 2.50 | -1.37 |
| UBA1 | Ubiquitin-like modifier-activating enzyme 1 | 2.50 | -1.32 |
| UBE2G1 | Ubiquitin-conjugating enzyme E2 G1; Ubiquitin-conjugating enzyme E2 G1, N-terminally processed | 2.43 | -1.52 |
| CTDSP1 | Carboxy-terminal domain RNA polymerase II polypeptide A small phosphatase 1; CTD small phosphatase-like protein | 2.32 | -1.59 |
| NFE2L1 | Nuclear factor erythroid 2-related factor 1 | 2.27 | -1.76 |
| DNAJB12 | DnaJ homolog subfamily B member 12 | 2.26 | -1.60 |
| UBE2J1 | Ubiquitin-conjugating enzyme E2 J1 | 2.15 | -1.59 |
| UBE2C | Ubiquitin-conjugating enzyme E2 C | 2.12 | -1.37 |
| UBE3C | Ubiquitin-protein ligase E3C | 2.01 | -1.68 |
| VPRBP (DCAF1) | DDB1 And CUL4 Associated Factor 1 | 2.00 | -1.69 |
| LSR | Lipolysis-stimulated lipoprotein receptor | 1.99 | -2.23 |
| UBA6 | Ubiquitin-like modifier-activating enzyme 6 | 1.86 | -1.04 |
| DIABLO | Diablo homolog, mitochondrial | 1.83 | -2.10 |
| KDM5B | Lysine-specific demethylase 5B | 1.82 | -1.16 |
| NEDD4L | E3 ubiquitin-protein ligase NEDD4-like | 1.74 | -1.35 |
| PRIM1 | DNA primase small subunit | 1.73 | -1.26 |
| CWC27 | Peptidyl-prolyl cis-trans isomerase CWC27 homolog | 1.73 | -1.00 |
| PDLIM5 | PDZ and LIM domain protein 5 | 1.70 | -1.42 |
| C15orf39 | Uncharacterized protein C15orf39 | 1.70 | -1.59 |
| YEATS2 | YEATS domain-containing protein 2 | 1.66 | -1.65 |
| XIAP | E3 ubiquitin-protein ligase XIAP | 1.65 | -1.10 |
| PAIP1 | Polyadenylate-binding protein-interacting protein 1 | 1.63 | -1.72 |
| BCOR | BCL-6 corepressor | 1.60 | -2.25 |
| DCTPP1 | dCTP pyrophosphatase 1 | 1.60 | -1.10 |
| CDC34 | Ubiquitin-conjugating enzyme E2 R1 | 1.58 | -1.98 |
| SNX7 | Sorting nexin-7 | 1.57 | -1.75 |
| SUGP2 | SURP and G-patch domain-containing protein 2 | 1.47 | -1.80 |
| EMD | Emerin | 1.45 | -1.34 |
| CDKN2A | Tumor suppressor ARF | 1.44 | -1.07 |
| TTK | Dual specificity protein kinase TTK | 1.42 | -2.99 |
| NUP54 | Nucleoporin p54 | 1.41 | -1.06 |
| NUDT21 | Cleavage and polyadenylation specificity factor subunit 5 | 1.39 | -1.53 |
| EEF1A2 | Elongation factor 1-alpha 2 | 1.38 | -1.19 |
| TUBGCP2 | Gamma-tubulin complex component 2 | 1.37 | -1.11 |
| UBE2R2 | Ubiquitin-conjugating enzyme E2 R2 | 1.37 | -1.31 |
| GTF3C2 | General transcription factor 3C polypeptide 2 | 1.36 | -1.60 |
| XRN2 | 5-3 exoribonuclease 2 | 1.36 | -1.53 |
| DYNC1LI1 | Cytoplasmic dynein 1 light intermediate chain 1 | 1.34 | -1.52 |
| RPN2 | Dolichyl-diphosphooligosaccharide--protein glycosyltransferase subunit 2 | 1.33 | -1.29 |
| HNRNPA3 | Heterogeneous nuclear ribonucleoprotein A3 | 1.32 | -1.38 |
| UBE2D1 | Ubiquitin-conjugating enzyme E2 D1 | 1.32 | -1.52 |
| UBR5 | E3 ubiquitin-protein ligase UBR5 | 1.31 | -1.52 |
| FLNC | Filamin-C | 1.31 | -1.18 |

**Supplementary Table 2.** Ubiquitylated proteins up-regulated with HERC1 WT vs TBL.

| Gene name | Protein description | p-value (-log<sub>10</sub>) | WT/TBL ratio (log<sub>2</sub>) |
|---|---|---|---|
| ANXA5 | Annexin A5 | 4.37 | 1.62 |
| TCEAL4 | Transcription elongation factor A protein-like 4; Transcription elongation factor A protein-like 2 | 3.54 | 1.66 |
| EIF2S3 | Eukaryotic translation initiation factor 2 subunit 3; Putative eukaryotic translation initiation factor 2 subunit 3-like protein | 3.42 | 1.25 |
| ABHD17B | Alpha/beta hydrolase domain-containing protein 17B | 3.08 | 1.77 |
| WDR43 | WD repeat-containing protein 43 | 3.04 | 1.45 |
| AKAP9 | A-kinase anchor protein 9 | 2.83 | 1.05 |
| USP6NL | USP6 N-terminal-like protein | 2.75 | 2.13 |
| ASNS | Asparagine synthetase [glutamine-hydrolyzing] | 2.66 | 1.98 |
| PSMC4 | 26S protease regulatory subunit 6B | 2.51 | 1.12 |
| AFF4 | AF4/FMR2 family member 4 | 2.50 | 1.55 |
| SYNE2 | Nesprin-2 | 2.45 | 2.65 |
| TMPO | Lamina-associated polypeptide 2, isoform alpha; Thymopoietin; Thymopentin | 2.33 | 1.53 |
| PSMC6 | 26S protease regulatory subunit 10B | 2.31 | 2.73 |
| VAMP8 | Vesicle-associated membrane protein 8 | 2.30 | 1.95 |
| DHPS | Deoxyhypusine synthase | 2.27 | 2.06 |
| SF1 | Splicing factor 1 | 2.27 | 1.94 |
| GNE | Bifunctional UDP-N-acetylglucosamine 2-epimerase/N-acetylmannosamine kinase; UDP-N-acetylglucosamine 2-epimerase (hydrolyzing); N-acetylmannosamine kinase | 2.21 | 1.44 |
| CXADR | Coxsackievirus and adenovirus receptor | 2.14 | 1.41 |
| EXOC6 | Exocyst complex component 6 | 2.14 | 1.55 |
| SUMO3 | Small ubiquitin-related modifier 3; Small ubiquitin-related modifier 4 | 1.97 | 1.52 |
| PIK3CA | Phosphatidylinositol 4,5-bisphosphate 3-kinase catalytic subunit alpha isoform | 1.87 | 1.35 |
| EIF2S2 | Eukaryotic translation initiation factor 2 subunit 2 | 1.78 | 1.32 |
| VAC14 | Protein VAC14 homolog | 1.71 | 1.55 |
| CDC123 | Cell division cycle protein 123 homolog | 1.56 | 1.04 |
| CTBP1 | C-terminal-binding protein 1 | 1.50 | 1.64 |
| DAAM1 | Disheveled-associated activator of morphogenesis 1 | 1.45 | 1.66 |
| NR1D2 | Nuclear receptor subfamily 1 group D member 2 | 1.45 | 1.00 |
| THADA | Thyroid adenoma-associated protein | 1.44 | 1.15 |
| CORO1C | Coronin-1C | 1.42 | 1.48 |
| UBA2 | SUMO-activating enzyme subunit 2 | 1.41 | 2.49 |
| AUP1 | Ancient ubiquitous protein 1 | 1.40 | 1.23 |
| CIAPIN1 | Anamorsin | 1.38 | 1.25 |
| PAPSS1 | Bifunctional 3-phosphoadenosine 5-phosphosulfate synthase 1; Sulfate adenylyltransferase; Adenylyl-sulfate kinase | 1.38 | 1.58 |
| NUP205 | Nuclear pore complex protein Nup205 | 1.37 | 1.45 |
| SBSN | Suprabasin | 1.35 | 1.42 |
| ZSWIM6 | Zinc finger SWIM domain-containing protein 6 | 1.33 | 1.63 |
| HIST1H2AC(H2AC6) | Histone H2A type 1-C; Histone H2A type 3; Histone H2A type 1-B/E | 1.32 | 1.02 |

**Supplementary table 3.** Ubiquitylated proteins down-regulated with HERC1 WT vs TBL.

| **Gene name** | **Protein description** | **p-value<br>(-log<sub>10</sub>)** | **WT/TBL ratio<br>(log<sub>2</sub>)** |
|---|---|---|---|
| TIMM13 | Mitochondrial import inner membrane translocase subunit Tim13 | 3.55 | -4.07 |
| UBA1 | Ubiquitin-like modifier-activating enzyme 1 | 3.28 | -1.89 |
| UBE2J1 | Ubiquitin-conjugating enzyme E2 J1 | 3.23 | -1.85 |
| CS | Citrate synthase, mitochondrial | 2.82 | -6.21 |
| TANK | TRAF family member-associated NF-kappa-B activator | 2.65 | -3.46 |
| UBE2C | Ubiquitin-conjugating enzyme E2 C | 2.63 | -1.33 |
| UBE2N | Ubiquitin-conjugating enzyme E2 N; Putative ubiquitin-conjugating enzyme E2 N-like | 2.36 | -1.44 |
| UBE2G1 | Ubiquitin-conjugating enzyme E2 G1; Ubiquitin-conjugating enzyme E2 G1, N-terminally processed | 2.33 | -1.51 |
| UBE3C | Ubiquitin-protein ligase E3C | 2.12 | -1.35 |
| SLX4 | Structure-specific endonuclease subunit SLX4 | 2.06 | -1.77 |
| DIABLO | Diablo homolog, mitochondrial | 2.01 | -2.24 |
| ZC3H15 | Zinc finger CCCH domain-containing protein 15 | 1.91 | -2.64 |
| UBE2L3 | Ubiquitin-conjugating enzyme E2 L3 | 1.89 | -2.16 |
| PPM1B | Protein phosphatase 1B | 1.88 | -2.07 |
| NEDD4L | E3 ubiquitin-protein ligase NEDD4-like | 1.88 | -1.47 |
| LSR | Lipolysis-stimulated lipoprotein receptor | 1.87 | -1.97 |
| PDLIM5 | PDZ and LIM domain protein 5 | 1.67 | -1.40 |
| SNX7 | Sorting nexin-7 | 1.67 | -1.85 |
| BCOR | BCL-6 corepressor | 1.65 | -2.23 |
| XIAP | E3 ubiquitin-protein ligase XIAP | 1.63 | -1.08 |
| HIST1H2AA | Histone H2A type 1-A; Histone H2AX | 1.58 | -2.45 |
| CPNE1 | Copine-1 | 1.56 | -2.00 |
| ATP6V1C1 | V-type proton ATPase subunit C 1 | 1.52 | -1.62 |
| CAP1 | Adenylyl cyclase-associated protein 1 | 1.50 | -1.81 |
| CPSF1 | Cleavage and polyadenylation specificity factor subunit 1 | 1.49 | -1.02 |
| XRCC1 | DNA repair protein XRCC1 | 1.47 | -1.21 |
| HMGCS1 | Hydroxymethylglutaryl-CoA synthase, cytoplasmic | 1.47 | -1.18 |
| EMD | Emerin | 1.46 | -1.06 |
| CRTAP | Cartilage-associated protein | 1.46 | -1.85 |
| RCN2 | Reticulocalbin-2 | 1.45 | -1.09 |
| GTF3C2 | General transcription factor 3C polypeptide 2 | 1.42 | -1.31 |
| UBE2D1 | Ubiquitin-conjugating enzyme E2 D1 | 1.41 | -1.47 |
| CDC34 | Ubiquitin-conjugating enzyme E2 R1 | 1.40 | -1.60 |
| PPP2R5E | Serine/threonine-protein phosphatase 2A 56 kDa regulatory subunit epsilon isoform | 1.38 | -1.37 |
| UBE2Q1 | Ubiquitin-conjugating enzyme E2 Q1 | 1.38 | -2.54 |
| IL17RA | Interleukin-17 receptor A | 1.37 | -1.17 |
| OXSR1 | Serine/threonine-protein kinase OSR1 | 1.36 | -1.59 |
| BIRC2 | Baculoviral IAP repeat-containing protein 2 | 1.31 | -1.91 |

